# Visual modulation of spectrotemporal receptive fields in mouse auditory cortex

**DOI:** 10.1101/2021.08.06.455445

**Authors:** James Bigelow, Ryan J. Morrill, Timothy Olsen, Stephanie N. Bazarini, Andrea R. Hasenstaub

**Affiliations:** Coleman Memorial Laboratory; Neuroscience Graduate Program; Department of Otolaryngology–Head and Neck Surgery, University of California, San Francisco, 94143

## Abstract

Recent studies have established significant anatomical and functional connections between visual areas and primary auditory cortex (A1), which may be important for perceptual processes such as communication and spatial perception. However, much remains unknown about the microcircuit structure of these interactions, including how visual context may affect different cell types across cortical layers, each with diverse responses to sound. The present study examined activity in putative excitatory and inhibitory neurons across cortical layers of A1 in awake male and female mice during auditory, visual, and audiovisual stimulation. We observed a subpopulation of A1 neurons responsive to visual stimuli alone, which were overwhelmingly found in the deep cortical layers and included both excitatory and inhibitory cells. Other neurons for which responses to sound were modulated by visual context were similarly excitatory or inhibitory but were less concentrated within the deepest cortical layers. Important distinctions in visual context sensitivity were observed among different spike rate and timing responses to sound. Spike rate responses were themselves heterogeneous, with stronger responses evoked by sound alone at stimulus onset, but greater sensitivity to visual context by sustained firing activity following transient onset responses. Minimal overlap was observed between units with visual-modulated firing rate responses and spectrotemporal receptive fields (STRFs) which are sensitive to both spike rate and timing changes. Together, our results suggest visual information in A1 is predominantly carried by deep layer inputs and influences sound encoding across cortical layers, and that these influences independently impact qualitatively distinct responses to sound.

**Significance statement:** Multisensory integration is ubiquitous throughout the brain, including primary sensory cortices. The present study examined visual responses in primary auditory cortex, which were found in both putative excitatory and inhibitory neurons and concentrated in the deep cortical layers. Visual-modulated responses to sound were similarly observed in excitatory and inhibitory neurons but were more evenly distributed throughout cortical layers. Visual modulation moreover differed substantially across distinct sound response types. Transient stimulus onset spike rate changes were far less sensitive to visual context than sustained spike rate changes during the remainder of the stimulus. Spike timing changes were often modulated independently of spike rate changes. Audiovisual integration in auditory cortex is thus diversely expressed among cell types, cortical layers, and response types.

## Introduction

Evidence accumulated within recent decades demonstrates that primary auditory cortex (A1) is not exclusively involved in processing sound. Instead, A1 integrates information carried by projections from multiple sensory and motor areas with input from the ascending auditory pathway (Schneider and Mooney, 2018; King et al., 2019). For many species including humans, monkeys, and mice, visual projections comprise a particularly dense source of input to A1 (Banks et al., 2011). This likely reflects the tendency of environmental features and events to be simultaneously transduced by the auditory and visual modalities, thus giving rise to audiovisual perceptual processes such as spatial localization and communication. Consistent with these observations, physiological studies have found that sound-evoked responses may be modulated by simultaneously presented visual stimuli, either increasing or decreasing firing rates relative to sound alone (Bizley and King, 2008; Kayser et al., 2009).

Most physiological studies of audiovisual integration in A1 have not investigated potential differences among cortical layers and cell types (e.g., excitatory vs. inhibitory). This is surprising, as an essential aspect of cortical organization is its division into layers, each with distinct cell type compositions and connectivity patterns with cortical and subcortical structures. Consistent with these anatomical differences, numerous studies in A1 have reported differences in sound encoding properties of neurons across cortical layers (Atencio et al., 2009) and cell types (Atencio and Schreiner, 2008; Phillips et al., 2017b). These findings raise the possibility that multisensory integrative properties of A1 might similarly vary by cortical layer and cell type. Indeed, a recent study from our lab found that a subset of neurons in mouse A1 were responsive to visual flash stimuli, and that these neurons were concentrated in the infragranular layers (Morrill and Hasenstaub 2018). However, this study left open the question of whether neurons with audiovisual integrative responses, such as visual-modulated responses to sound or responses to both modalities, are distributed in parallel with the infragranular visual-responsive neurons. Similarly, whether unimodal visual responses or audiovisual integrative responses differ between neuron types (excitatory, inhibitory) remains to be investigated.

With few exceptions (Kayser et al., 2010; Atilgan et al., 2018), studies examining multisensory integration in A1 and elsewhere have relied on changes in time-averaged spike rates to quantify integrative effects. However, neurons throughout the auditory pathway may encode sound features and other events through changes in spike timing, rate, or both (deCharms and Merzenich 1996; Malone et al., 2010; Insanally et al., 2019). Capturing both spike rate and timing changes in A1 may be fundamental to understanding audiovisual integration for two reasons. First, neurons with spike timing changes alone are common in A1, in some preparations reflecting the majority (Insanally et al., 2019). Thus, focusing on spike-rate changes alone may underestimate the prevalence or strength of multisensory integrative activity. Second, downstream targets of A1 may be differently influenced by spike rate and timing changes. Capturing spike timing effects may therefore provide insight into multisensory processes in structures receiving projections from A1.

Spike-rate changes are themselves multifaceted and may include transient firing changes at stimulus onset or offset, as well as sustained changes throughout the stimulus period (Lu et al., 2001; Wang et al., 2005; Malone et al., 2015). These diverse response types may reflect distinct network states (Churchland et al., 2010) and sources of information from the ascending auditory pathway (Liu et al., 2019). Resolving potential differences in multisensory integrative properties among response types may be similarly fundamental to understanding the nature and extent of multisensory integration in A1.

The current study examined single unit responses in awake mouse A1 to auditory, visual, and audiovisual stimulation. High-density multichannel electrode arrays enabled cortical depth estimation for each neuron and physiological features permitted classification of putative excitatory and inhibitory units. Broadband receptive field estimation stimuli delivered in segments enabled measurement of both transient onset and sustained firing rate responses, as well spectrotemporal receptive fields (STRFs), which are sensitive to both spike rate and timing changes.

## Materials and Methods

### Subjects and surgical preparation

All procedures were approved by the Institutional Animal Care and Use Committee at the University of California, San Francisco. A total of 15 adult mice (6 female) served as subjects (median age 99 days, range 58–169 days). All mice had a C57BL/6 background and expressed optogenetic effectors targeting interneuron subpopulations, which were not manipulated in the current experiment. Mice were housed in groups of two to five under a 12H-12H light-dark cycle. Surgical procedures were performed under isoflurane anesthesia with perioperative analgesics (lidocaine, meloxicam, and buprenorphine) and monitoring. A custom stainless steel headbar was affixed to the cranium above the right temporal lobe with dental cement, after which subjects were allowed to recover for at least two days. Prior to electrophysiological recording, a small craniotomy (~1–2 mm diameter) centered above auditory cortex (~2.5–3.5 mm posterior to bregma and under the squamosal ridge) was made within a window opening in the headbar. The craniotomy was then sealed with silicone elastomer (Kwik-Cast, World Precision Instruments). The animal was observed until ambulatory (~5–10 min) and allowed to recover for a minimum of 2 h prior to electrophysiological recording. The craniotomy was again sealed with silicone elastomer at the conclusion of recording, and the animal was housed alone thereafter. Electrophysiological recordings were conducted for each animal on up to five consecutive days following the initial craniotomy procedure.

### Auditory and visual stimuli

All stimuli were generated in MATLAB (Mathworks) and delivered using Psychophysics Toolbox Version 3 (Kleiner et al., 2007). Sounds were delivered through a free-field electrostatic speaker (ES1, Tucker-Davis Technologies) approximately 15–20 cm from the left (contralateral) ear using an external soundcard (Quad Capture, Roland) at a sample rate of 192 kHz. Sound levels were calibrated to 60 ± 5 dB at ear position (Model 2209 meter, Model 4939 microphone, Brüel & Kjær). Visual stimuli were presented on a 19-inch LCD monitor with a 60 Hz refresh rate (ASUS VW199 or Dell P2016t) centered 25 cm in front of the mouse. Monitor luminance was calibrated to 25 cd/m^2^ for 50% gray at eye position.

For the majority of recordings, search stimuli used for cortical depth estimation included click trains, noise bursts, and pure tone pips, plus the experimental stimuli described below. For a small minority of recordings, only tone pips and experimental stimuli were presented due to time constraints. In some recordings, additional search stimuli were presented, such as frequency-modulated sweeps. Click trains comprised broadband 5 ms non-ramped white noise pulses presented at 4 Hz for 1 s at 60 dB with a ~1 s interstimulus interval (ISI), with 20–50 repetitions. Noise bursts consisted of 50 ms non-ramped band-passed noise with a uniform spectral distribution between 4 and 64 kHz presented at 60 dB in 500 unique trials with a ~350 ms ISI. Pure tones consisted of 100 ms sinusoids with 5-ms cosine-squared onset/offset ramps presented at a range of frequencies (4–64 kHz, 0.2 octave spacing) and attenuation levels (30– 60 dB, 5 dB steps). Three repetitions of each frequency-attenuation combination were presented in pseudorandom order with an ISI of ~550 ms. Peristimulus-time histograms (PSTHs) quantifying time-binned multi-unit firing rates were constructed for each stimulus. For tone pips, frequency-response area (FRA) functions were constructed from baseline-subtracted spike counts during the stimulus period averaged across trials at each frequency-attenuation combination. PSTHs and FRAs from an example recording are shown in **Figure 1C**.

**Figure 1.**
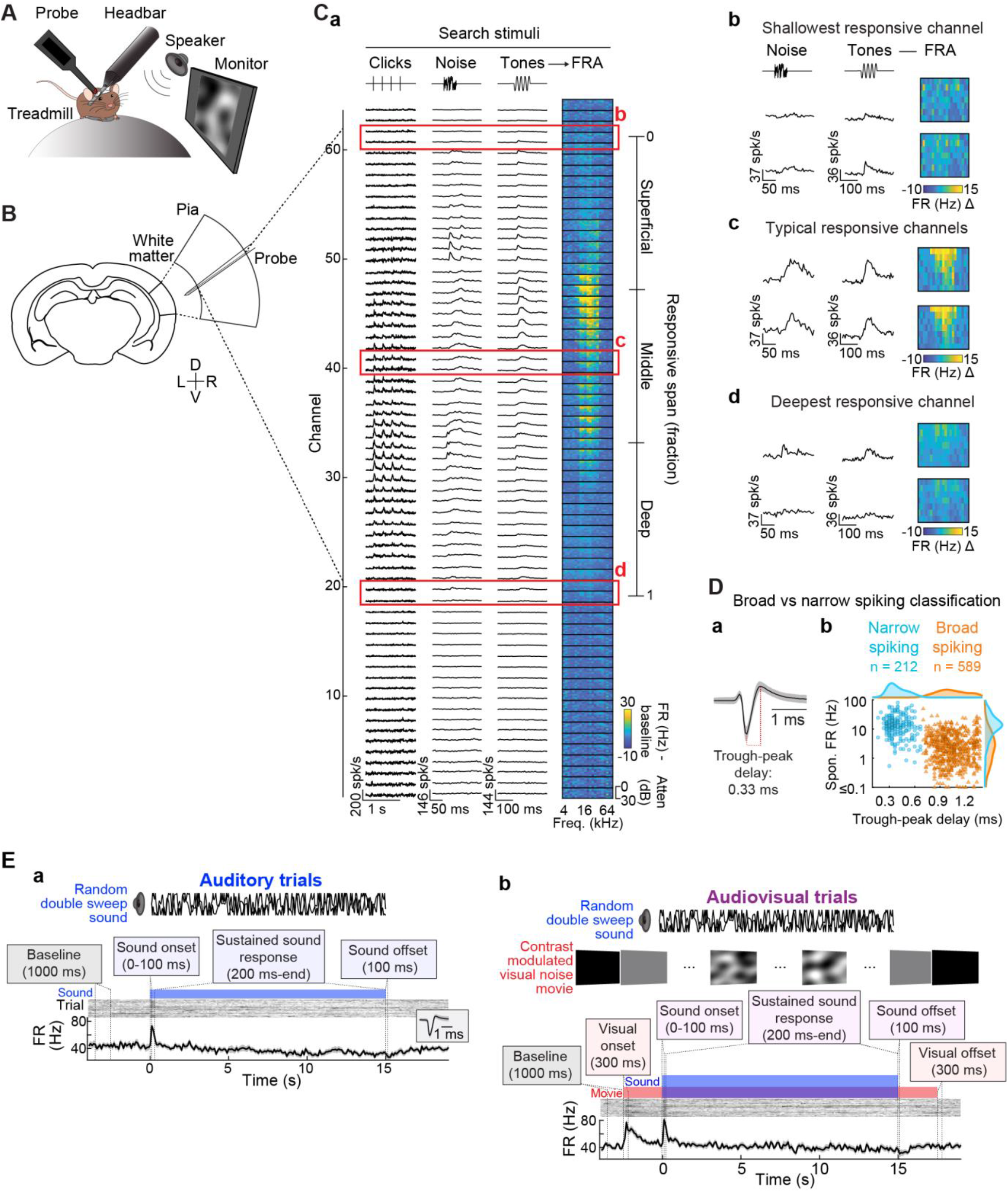
Single unit recording and audiovisual stimulation in awake mouse auditory cortex. (**A**) Mice were head fixed atop a spherical treadmill. A headbar window provided access to primary auditory cortex (A1) of the right hemisphere for extracellular recording with translaminar probes. Sounds were presented to the contralateral ear through an electrostatic speaker and visual stimuli were presented via a monitor centered in front of subjects at 25 cm distance. (**B**) Coronal mouse brain section with magnification of A1 and linear multichannel electrode arrays (64-channels, 20μm spacing) used to simultaneously record neuronal activity across all cortical layers. (**C**) Auditory cortical depth estimation. (**a**) Multiunit responses evoked by search stimuli (e.g., click trains, noise bursts, pure tones) were used to guide visual demarcation of the span of responsive channels which served as an estimate of the putative cortical span and was used to assign a fractional depth value to each recorded neuron. Fractional responsive span was further divided into Superficial, Middle, and Deep bins. (**b–d**) Example multiunit responses from (b) the shallowest channel of the responsive span, (c) responsive channels from the middle of the probe, and (d) deepest channel of the responsive span. (**D**) Identification of putative excitatory and inhibitory neurons by waveform morphology clustering. (**a**) Example single-unit waveform (black line: median, gray shading: median absolute deviation) showing trough-peak delay calculation. (**b**) The distribution of trough-peak delay times was sharply bimodal, permitting straightforward identification of broad spiking (BS; putative excitatory) and narrow spiking (NS; putative inhibitory) neurons. BS and NS unit populations were further distinguished by differences in spontaneous firing rate. (**E**) Auditory and visual stimulation paradigm. (**a**) Auditory trials comprised non-repeating 15-s segments of a random double sweep (RDS) stimulus, comprising two continuously frequency-modulated pure tones which varied independently of one another between 4 and 64 kHz. Trials were separated by silent intertrial intervals (~4–9 s), permitting calculation of spontaneous firing rates. Sound onset firing rate responses were defined by a 100-ms window post-stimulus onset. Sustained firing rates were quantified within a window 200-ms post-stimulus onset to the end of the stimulus (15 s). Inset shows the example unit spike waveform (median ± MAD). (**b**) Audiovisual trials were identical to Auditory trials (including the same RDS segments) with the addition of a visual contrast modulated noise (CMN) stimulus. The CMN stimulus led and trailed the RDS stimulus by 2.5 s, permitting unambiguous assessment of visual onset and offset firing rate responses and allowing adaptation to the visual stimulus prior to sound onset. The auditory (RDS) and visual (CMN) stimuli were uncorrelated with each other. Auditory and Audiovisual trials were interleaved in pseudorandom order.

As depicted by **Figure 1E**, experiments comprised two trial types: (a) Auditory trials, which presented sound only, and (b) Audiovisual trials, which included both sound and visual stimuli. For both trial types, the auditory stimulus was a random double sweep (RDS), a continuous, spectrally sparse receptive field estimation stimulus capable of effectively driving activity across diversely tuned neurons in A1 (Gourévitch et al., 2015). The RDS comprised two uncorrelated random sweeps, each varying continuously and smoothly over time between 4 and 64 kHz, with a maximum sweep modulation frequency of 20 Hz. Sample RDS frequency vectors are depicted in **Figure 1E** and **Figure 4A, a**. The RDS was delivered in 15-s non-repeating segments (40 trials, 10 minutes total stimulation; cf. Rutkowski et al., 2002). The inter-sound interval was ~9 s, with visual stimuli trailing and leading sounds within this interval. Thus, intertrial intervals were ~9 s for consecutive auditory trials, ~4 s for consecutive audiovisual trials, and ~6.5 seconds for mixed trial type sequences. The same 40 RDS segments were used for Auditory and Audiovisual trials to maintain identical stimulus statistics between conditions. Audiovisual trials were thus identical to Auditory trials with the exception of an additional contrast modulated visual noise stimulus.

As described in detail elsewhere (Niell and Stryker, 2008), CMN is a broadband stimulus designed to drive as many primary visual cortical neurons as possible. The stimulus is generated by first creating a random frequency spectrum in the Fourier domain. The temporal frequency spectrum was flat with a low-pass cutoff at 10 Hz. The spatial frequency spectrum dropped off as A(f) ~ 1/(f+f_c_), with f_c_ = 0.05 cycles/°. A spatiotemporal movie was then created by inverting the three-dimensional spectrum. Finally, contrast modulation was imposed by multiplying the movie by a sinusoidally variable contrast function. The CMN stimulus was generated at 60×60 pixels, then interpolated to 900×900 pixels. The first and last frames of the CMN movie were uniform 50% gray, providing abrupt luminance changes at stimulus onset and offset from black during the intertrial interval. The CMN stimulus led and trailed the RDS sounds by 2.5 s to allow ample time for potential visual-evoked spiking responses to reach an adapted state prior to sound onset responses and persist throughout sound offset responses.

### Electrophysiology

Recordings were conducted inside a sound attenuation chamber (Industrial Acoustics Company). Anesthesia has well known and profound influences on auditory cortical encoding, including the relative prevalence of onset and sustained firing rate responses (Wang et al., 2005). Recordings were thus conducted in awake, headfixed animals moving freely atop a spherical treadmill in **Figure 1A** (Dombeck et al., 2007; Niell and Stryker, 2010; Phillips and Hasenstaub, 2016; Phillips et al., 2017a, 2017b; Morrill and Hasenstaub, 2018; Bigelow et al., 2019). The silicone elastomer filling the craniotomy was removed and a single shank, linear multichannel electrode array (Cambridge Neurotech) was slowly lowered into cortex using a motorized microdrive (FHC). Arrays with 64 channels (20 μm site spacing, 1260 μm total span) were used for all recordings except one, which used a 32-channel array (25 μm site spacing, 775 μm total span). Prior to lowering the probe, the craniotomy was filled with 2% agarose to stabilize the brain surface. After reaching depths of approximately 800–1000 μm below the first observation of action potentials, probes were allowed to settle for at least 20 mins before initiating recording. Continuous extracellular voltage traces were collected using an RHD2000 (Intan Technologies) at a sample rate of 30 kHz. Other experimental events such as stimulus event times were stored concurrently by the same system.

The span of recording channels (1260 μm) exceeded mouse cortical depth (~800 μm; Paxinos and Franklin, 2019), resulting in a majority of sound-responsive channels plus an additional subset of channels recorded outside of A1. As depicted in **Figure 1C**, multi-unit responses evoked by search stimuli (e.g., click trains, noise bursts, tone pips) were used to guide visual demarcation of the range of sound-responsive channels, which served as an estimate of cortical span. Although penetrations were approximately perpendicular to the cortical surface, it was impractical to achieve perfect orthogonality. Thus, the responsive span of each recording was normalized such that each channel was expressed as a fraction of total depth (Morrill and Hasenstaub, 2018). We note that this cortical depth estimation procedure is less precise than our prior study, in which Di-I was applied to the probe for histologically referenced depth estimation (Morrill and Hasenstaub, 2018). Nevertheless, we observed parallel depth distributions of visual responsive neurons in the current and prior studies, suggesting the current method achieved a rough approximation to the histological approach. However, due to the lack of histological verification, a more conservative depth categorization approach was adopted, dividing the responsive span into three equal bins reflecting superficial, middle, and deep-layer neuron populations.

Recordings targeted A1 using stereotaxic coordinates and anatomical landmarks such as characteristic vasculature patterns (Joachimsthaler et al., 2014). Previous studies have reported significant differences in tone onset latencies between primary and non-primary auditory cortical fields (Joachimsthaler et al., 2014), with latencies between 5 and 18 ms for primary fields (median ~9 ms), and 8–32 ms for non-primary fields (median ~12–16 ms). Thus, tone onset latencies were used to support designations of putative primary recording sites. PSTHs were constructed from multi-unit activity (negative threshold crossings exceeding 4.5 median absolute deviations of the continuous voltage trace distribution) using 2-ms bins and smoothed with a Savitzky-Golay filter (3rd order, 10-ms window). Onset latency was defined as the first post-stimulus bin in which the firing rate exceeded 2.5 standard deviations of the pre-stimulus firing rate bins (Morrill and Hasenstaub, 2018). Recordings for which the median latency across the responsive channel span was 14 ms or less were considered putative primary sites and retained for further analysis (49 of 60 total recordings). The retained putative primary recordings universally exhibited robust multi-unit responses to click trains, noise bursts, and tone pips, and clear evidence of frequently-level tuning in the FRA plots (**Figure 1C**) as well as spectrotemporal tuning in single-unit responses to RDS stimuli (**Figure 4**).

Single-unit activity was isolated from continuous multichannel traces using Kilosort 2.0 (Pachitariu et al., 2016; available: https://github.com/MouseLand/Kilosort) and further validated by auto- and cross-correlation analysis, refractory period analysis, and cluster isolation statistics. Although the majority of isolated units were held throughout the entire recording (~28 minutes), isolation of individual neurons was occasionally disrupted and lost partway through the experiment. Thus, the active timespan for each unit was estimated by visual demarcation of unit activity plots over time. Inactive trials were discarded from further analysis. For the remaining subset of active trials, RDS stimuli were matched between conditions by only using available RDS segments common to both conditions. This ensured strict equivalence of auditory stimuli between conditions, isolating any observed differences to the presence of the visual CMN stimulus. Only units with 10 or more active trials (5 per condition) were retained for final analysis. A total of 801 units were included in the analyses below. As in previous publications (Phillips et al., 2017; Bigelow et al., 2019), units were classified as narrow-spiking (NS; expected to be overwhelmingly inhibitory) or broad-spiking (BS; expected to be mainly excitatory) on the basis of a clear bimodal distribution of waveform trough-peak delays (**Figure 1D**; NS, <600 μs, n = 212; BS, ≥600 μs, n = 589). Consistent with this classification, NS units had characteristically higher spontaneous firing rates than BS units (estimated from baseline period shown in **Figure 1E, a**; F = 391.31, p < 10^−70^, *η*^2^ = 0.329).

### Spectrotemporal receptive field estimation

Spectrotemporal fields (STRFs) were estimated using standard reverse-correlation techniques (Wu et al., 2006) as depicted in **Figure 4A, a–c**. RDS stimuli were discretized in 1/8 oct frequency bins and 5-ms time bins, which is sufficient resolution for modeling response properties in the majority of A1 neurons (Thorson et al., 2015). The spike-triggered average (STA) was obtained by adding the discretized stimulus segment preceding each spike to a cumulative total, and then dividing by the total spike count. For all data analyses, the peri-spike time analysis window spanned 0–100 ms prior to spike event times, sufficient for capturing the full latency-adjusted temporal response periods of the majority of A1 neurons (Atencio and Schreiner 2013, See et al., 2018). A broader window was used for display purposes, spanning 200 ms before and 50 ms after spike event times. The 50 ms post-spike window was included as a visualization of estimated acausal values, i.e., those that would be expected by chance given the finite recording time, stimulus and spike timing statistics, and smoothing parameters (Gourévitch et al. 2015). The first 200 ms of the RDS response from each trial were dropped from all STA calculations analyses to minimize bias reflecting strong onset transients.

The STA thus reflects the average binned stimulus segment preceding spike events and can be viewed as a linear approximation to the optimal stimulus for driving neuronal firing (deCharms and Merzenich, 1998). As discussed by Rutkowski et al. (2002), the STA can be formalized as the probability (*P*) of a stimulus frequency *f* occurring at time *t*_i_-*τ* given that a spike occurred, as expressed by the equation

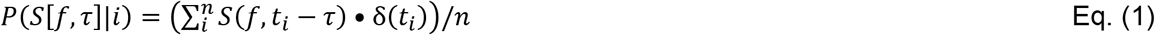

where *i* indicates a spike, *t*_i_ is a spiketime, *τ* is the time analysis window, *n* is the spike count, and Σ indicates summing across spikes. *S*(*f*,*t*) is the stimulus value at a given time-frequency bin, equaling one if an RDS frequency intersects the bin, two if both RDS frequencies coincide with the bin, and zero otherwise. *S*(*f*,*t*_i_-*τ*) represents the windowed stimulus aligned to a spike time. δ(*t*_i_) is equal to one if a spike occurs at time *t*_i_ and zero otherwise. With a slight modification, STRF time-frequency bins can be expressed in terms of deviation from mean driven firing rate (spikes/s - mean) using the terms defining the STA and Bayes’ theorem,

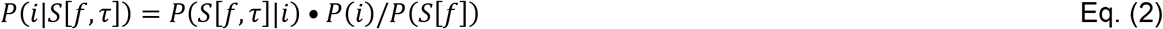

where *P*(*i*) is the probability of a spike occurring in a bin, equal to *n*_i_/*T*, where *T* is the total stimulus time and *P*(*S*[*f*]) is the probability of a tone frequency occurring in a bin. The mean driven firing rate is then subtracted from the STRF such that individual time-frequency bins reflect increases (positive, red) or decreases (negative, blue) from the mean driven rate (**Figure 4A, c**). Finally, the STRF is smoothed by a uniform 3×3 bin window to reduce overfitting to finite-sampled stimulus statistics. The STA and STRF expressed in units of spikes/s are multiples of each other since terms in the expression *P*(*i*)/*P*(*S*[*f*]) are constant and thus nearly identical for the purposes of all data analyses, including comparisons between conditions. However, we opt to report STRFs represented in firing-rate change units to facilitate interpretation of stimulus driven changes in neuronal activity.

In addition, ‘Null’ STRFs were calculated using identical procedures to those described above except that the stimulus was reversed in time, while preserving the original spike event times. This modification breaks the temporal relationship between the stimulus and spike times, but preserves spike count and timing statistics (e.g., interspike interval distribution), as well as the statistical distributions of the stimulus (Bigelow and Malone, 2017). The resulting STRF was used to estimate time-frequency bin values expected by chance within the constraints of finite spike counts and stimulus time.

STRFs were calculated independently for each condition. As in previous studies (Fritz et al., 2003), a difference STRF, which we refer to throughout as ΔSTRF, was calculated by subtracting the STRF obtained from Auditory trials from the STRF obtained from Audiovisual trials. A Null ΔSTRF was similarly obtained by subtracting Null Auditory STRF from the Null Audiovisual STRF.

### Mutual information analysis

Although the STRF effectively captures the spectrotemporal tuning of a neuron, it does not reveal the degree to which driven spiking activity is dependent upon similarity between the stimulus and receptive field. For instance, it is not apparent how consistently spiking activity is observed when the stimulus closely approximates the STRF (e.g., RDS frequency vectors intersecting STRF excitatory subfields) and whether spiking is inhibited when the stimulus is anticorrelated or uncorrelated with the STRF (e.g., RDS frequency vectors intersecting STRF inhibitory subfields or regions with values near zero). Thus, mutual information was calculated to quantify the relationship between probability of firing and similarity between the stimulus and STRF according to previously published methodology (Atencio et al., 2008; Atencio and Schreiner, 2016).

As depicted by **Figure 12A**, mutual information reflects a scaled ratio of two distributions: *P*(x), which reflects the ‘similarity’ between the STRF and all possible RDS segments in the experiment, and the other, *P*(x|spike), reflecting STRF-stimulus ‘similarity’ values at time bins in which a spike occurred. Stimulus-STRF ‘similarity’ (x) is operationally defined as the inner product (i.e., projection value) between the STRF and the stimulus segment of equivalent dimensions. Put another way, the STRF is convolved with the stimulus, yielding a projection value for each 5-ms time bin in the stimulus (**Figure 12A, a–c**). For ease of interpretation, raw projection values were standardized by subtracting the mean of the raw *P*(x) distribution and dividing by its standard deviation. Thus, highly positive and negative standardized values reflect stimulus segments highly similar and dissimilar to the STRF, respectively. Values near zero imply a random relationship between the stimulus and STRF. The continuously valued projection values were separated into nine linearly spaced bins (**Figure 12A, d**). As in prior studies, the most extreme positive bin was dropped from the information calculation due to undersampling (the most extreme STRF-stimulus matches are rare), resulting in eight analyzed bins. Finally, mutual information (bits/spike) is calculated by the calculating log2-transformed ratio of *P*(x|spike) divided by *P*(x), multiplying the result by *P*(x|spike), and summing across bins. In equation form,

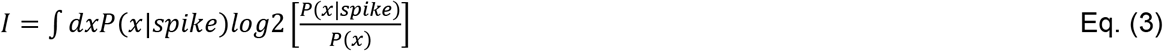

The information estimation procedure occasionally produced values of zero for a small minority of units, which were excluded from further information analyses. Information values were not calculated for units with <200 total spikes due to undersampling concerns. Intuitively, mutual information increases as the *P*(x) and *P*(x|spike) distributions diverge. For instance, if a given binned *P*(x) value is low but *P*(x|spike) is high, we conclude that the stimulus (and its associated ‘similarity’ value to the STRF) occurs infrequently but usually evokes a spiking response. Similarly, if *P*(x) is high but *P*(x|spike) is low, we infer that a regularly occurring stimulus typically inhibits spiking. In each scenario, the stimulus and response are thus mutually informative of one another. By contrast, if the binned *P*(x) and *P*(x|spike) values are similar (both high or low), the stimulus is not informative about spiking behavior.

### Statistical analysis

Our stimulus design allowed measurement of unimodal and bimodal evoked firing-rate responses as well as stimulus-driven changes in spike timing (e.g., spike alignment with RDS stimulus features). As indicated by **Figure 1E, a**, baseline (spontaneous) firing rates were measured from a window spanning 3.5 to 2.5 s prior to sound onset (immediately preceding visual stimulus onset for audiovisual trials). Sound onset responses were measured from spikes occurring within the first 100 ms of the stimulus. Sustained firing rates were estimated within the same window used for STRF estimation, from 200 ms post stimulus onset to the end of the stimulus (15 s). All sound-evoked firing rate response windows and STRF calculations were identical between conditions. Visual onset responses were analyzed in a window spanning 300 ms after the visual stimulus began. The wider window used to capture onset transients for visual compared to auditory stimuli accommodated the substantially longer response latencies typical of visual responses in A1 (Morrill and Hasenstaub, 2018).

Auditory and visual offset responses were similarly analyzed using 100- and 300-ms windows following stimulus offset, respectively. Unlike onset and sustained responses, offset responses required comparison against two baselines: first, the spontaneous window described above, and second, a window of equivalent duration (1 s) preceding the offset response (i.e., the final second of the stimulus). Differences from the second baseline ensured offset responses were not merely carryover from a sustained response. In addition, ‘significant’ offset responses were required to be above or below both baselines (but not between them) to ensure offset responses did not simply reflect the return of a sustained firing rate change to spontaneous activity. The larger of the two p-values was used to assess significance of the offset response, and its associated baseline was used for calculating effect size described below.

For each unit, the significance of all firing rate responses was assessed by comparison to baseline Wilcoxon signed-rank tests (paired) using **α** = 0.05. For all tests of individual unit significance, false discovery rate (FDR) was limited by implementing the Benjamini–Hochberg procedure with *q* = 0.05 across the unit population (Benjamini and Hochberg, 1995). The adjusted p-values produced by the FDR procedure are used to indicate significance for each response type for all example unit plots throughout the manuscript. For a standardized measure of response strength that accommodated both increases and decreases in firing rate from baseline, we defined effect sizes as the absolute difference between the evoked and spontaneous firing rate means, divided by the standard deviation of the spontaneous firing rate. This facilitated interpretation of response deviations from chance reflecting different analysis windows, unit types.

Significant differences in firing rate between conditions (auditory, audiovisual) were similarly assessed with Wilcoxon signed-rank tests (**α** = 0.05, Benjamini–Hochberg FDR correction with *q* = 0.05), and effect sizes were estimated as the absolute difference between conditions divided by the standard deviation of the auditory condition.

To assess the significance of STRFs, we used a reliability index in which the correlation coefficient was calculated between two STRFs computed from random trial halves (**Figure 4A, c**; Escabí et al., 2014). Reliability was defined as the mean across 1000 iterations for both ‘data’ (time-preserved stimulus) and ‘null’ (time-reversed stimulus) STRFs. A p-value was calculated reflecting the proportion of the null distribution exceeding the mean of the data distribution (**Figure 4A, e**). P-values equal to zero (cases where none of the null correlations exceeded the data mean) were adjusted to 0.000999 in reflection of the resolution permitted by the number of subsample iterations. Finally, the p-values were multiplied by two for an estimate of two-tailed significance and adjusted using Benjamini–Hochberg FDR correction (*q* = 0.05). Because the null subsampled STRF distribution skewed negative for some units (i.e., null reliability index > 0), we further required STRF reliability >0.2 in order to be considered ‘significant’. Similar to firing rate responses, STRF reliability effect sizes were estimated as the absolute difference between the data and null distributions, divided by the standard deviation of the null distribution. Significance of ΔSTRFs was assessed with the same approach, except using subsampled ΔSTRF correlation distributions (**Figure 9A, a–b**).

Except where otherwise noted, tests of population-level differences (e.g., among units in superficial, middle, deep cortical depth bins) were assessed by independent one-way analysis of variance (ANOVA), using effect sizes described above as the dependent variable. For uniformity in presenting the results, we use the same approach for testing between two variables (e.g., differences between NS and BS units), wherein ANOVA and the Student’s *t*-test produce equivalent p-values with F = *t*^2^.

## Results

The current study examined visual-modulation of sound-evoked responses in awake mouse A1. By delivering a continuous auditory receptive field estimation stimulus in 15-s segments separated by silent intervals, we were able to capture both transient onset firing rate responses as well as sustained firing rate responses throughout the duration of the stimulus. By using reverse correlation methods, we were further able to estimate STRFs, which are simultaneously sensitive to both spike rate and timing. Half of the trials included a continuous visual stimulus, enabling direct comparison of firing rate responses and STRFs between auditory alone and visual-modulated conditions. The visual stimulus both led and trailed the sound stimulus by 2.5 seconds, further allowing measurement of averaged firing rate responses evoked by the visual stimulus alone.

### Some A1 neurons respond to visual stimulation alone

Example visual-responsive units are shown in **Figure 2A–B**. Consistent with our earlier study examining visual-flash evoked responses in A1 (Morrill and Hasenstaub, 2018), we found that significant responses to the visual CMN stimulus were most prevalent in the deepest cortical depth bin (**Figure 2C**). Extending our previous work, we found that for a minority of units, visual responses reflected decreases in firing rate relative to baseline (e.g., **Figure 2B**). For a standardized measure of visual response strength that avoided overweighting units with high firing rates and accommodated both increases and decreases in firing rate from baseline, we defined effect sizes as the absolute difference between the evoked and spontaneous firing rate means, divided by the standard deviation of the spontaneous firing rate. One-way ANOVA confirmed that effect size of visual-evoked onset firing rate changes was significantly dependent upon cortical depth for both unit types, with the strongest responses in the deepest bin (NS: F = 13.95, p < 10^−5^, *η*^2^ = 0.118; BS: F = 7.62, p < 10^−3^, *η*^2^ = 0.025).

**Figure 2.**
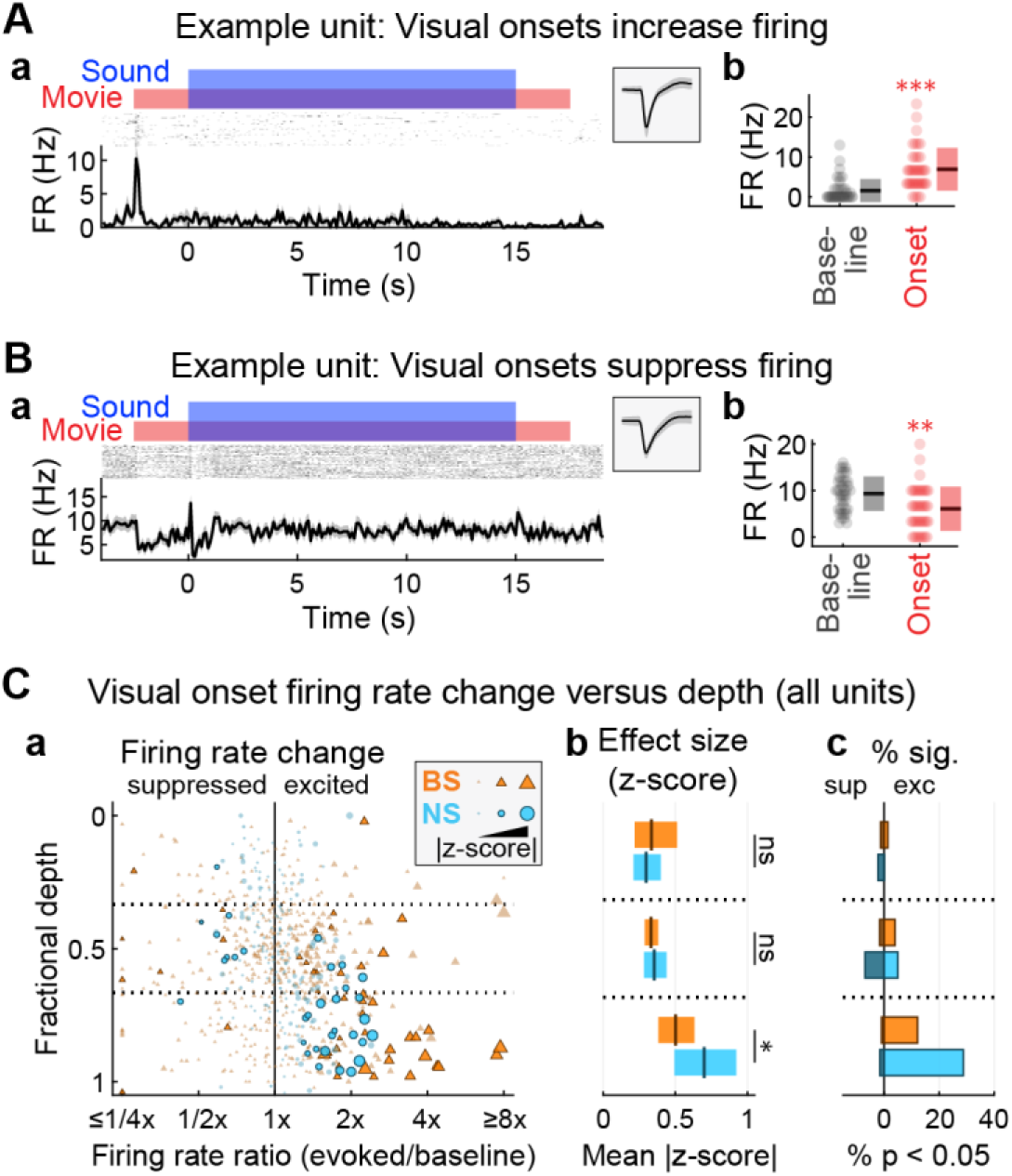
Some primary auditory cortical neurons respond to visual stimulation alone. (**A**) Example unit with an excitatory visual onset response. (**a**) Spiking responses quantified by peristimulus time histograms (lower) and binned-spike count matrices (upper) with blue and red bars indicating auditory and visual stimulus intervals, respectively (temporal binning: 100 ms). Inset shows the unit spike waveform (median ± MAD). (**b**) Summary of visual onset firing rate responses compared to baseline. Each dot represents mean firing rate for a single trial, with mean ± SD across trials indicated to the right. Wilcoxon signed-rank tests (paired): *p<0.05, **p<0.01, ***p<0.001, ns p>0.05. (**B**) Example unit with suppressed visual onset response. (**C**) Summary of visual onset firing rate responses by unit type and cortical depth. (**a**) Scatter plot depicting visual onset responses for each unit at its estimated fractional depth. Firing rate ratio values above and below 1 (x-axis) indicate excited and suppressed responses, respectively. Outlined markers indicate statistically significant responses (p < 0.05, Benjamini–Hochberg FDR correction). Marker sizes are scaled by effect size (absolute difference between onset and baseline means, divided by baseline SD). (**b**) Mean effect size (plus 99% confidence interval) across all recorded units (significant and non-significant visual onset responses included) by unit type and depth. Visual onset effect sizes for NS units were significantly stronger than BS units in the deepest cortical depth bin (but see Extended Data Figure 2-1 for spike-equated analysis). One-way ANOVA: *p<0.05, **p<0.01, ***p<0.001, ns p>0.05. (**c**) Histograms indicating percentages of all recorded units with significant excited and suppressed responses by unit type and depth. Significant visual responses were most concentrated in the deepest cortical depth bin.

Visual responses were detected in a larger percentage of NS units (15.6%) than BS units (7.8%), and visual responses were significantly stronger for NS units in the deepest cortical depth bin (F = 4.32, p = 0.039, *η*^2^ = 0.016). Differences were non-significant for the remaining depth bins (shallow: F = 0.25, p = 0.619, *η*^2^ = 0.002; middle: F = 0.29, p = 0.593, *η*^2^ = 0.001). We conducted follow-up analyses to test whether the larger percentage of visual responsive NS units could be explained by their characteristically higher firing rates – and thus statistical power for detecting firing rate changes. Mean firing rates were calculated for each unit across audiovisual trials (including the full stimulus period plus spontaneous activity 1 s before and after the stimulus), producing population mean rates of 18.11 Hz for NS units and 4.45 Hz for BS units (**Extended Data Figure 2-1A, a**). By randomly subsampling spikes from NS units with firing rates above the BS mean, we created a pseudo-population of NS units with mean firing rate equivalent to BS units (4.45 Hz; **Extended Data Figure 2-1A, b**). Visual onset response data were recalculated, including the p-value distribution which was readjusted by the Benjamini–Hochberg FDR procedure. As seen in **Extended Data Figure 2-1B**, differences in visual response effect sizes were non-significant for all depth bins (all F-ratios < 1.4, all p-values > 0.24). Thus, visual responses were observed in both NS and BS units, but any differences between these unit subpopulations may have reflected inherent differences in firing rate.

**Extended Data Figure 2-1.**
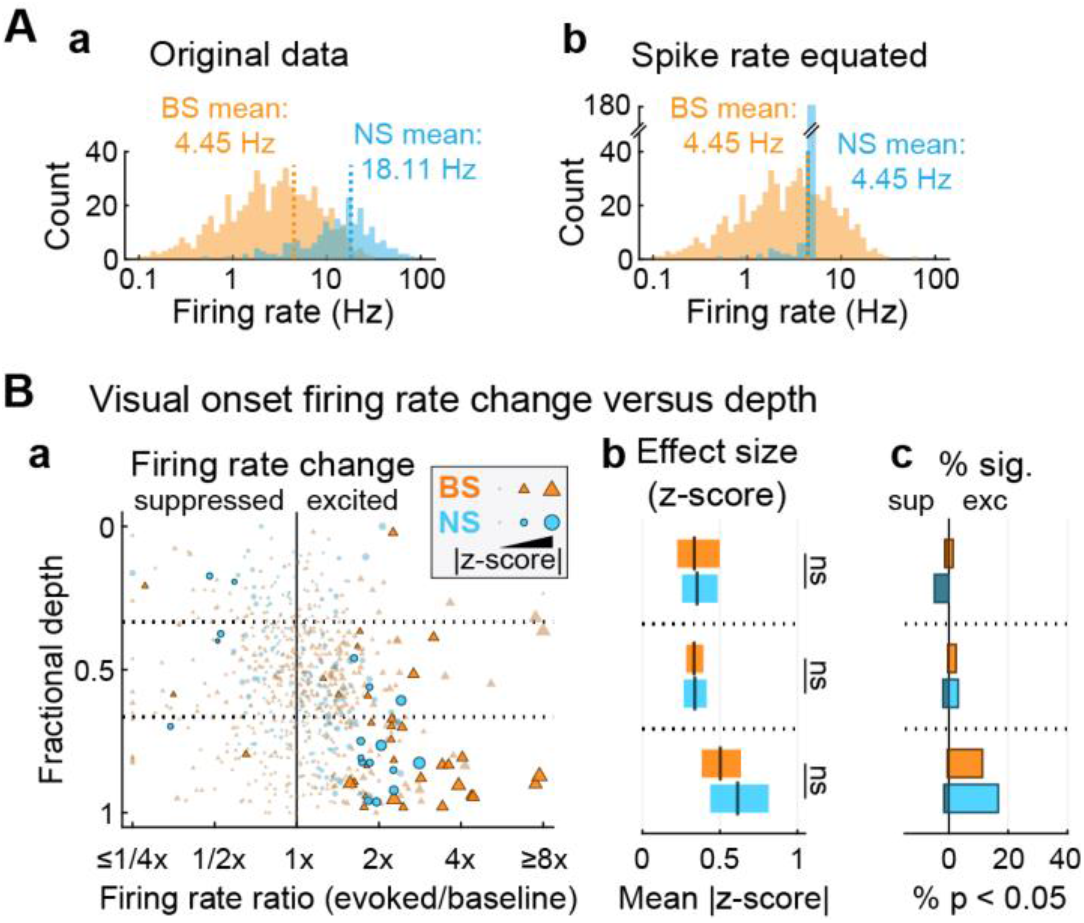
Differences in visual responses between NS and BS units may reflect inherent differences in firing rate. (**A**) Firing rate histograms for each unit type inclusive of both spontaneous and stimulated periods. (**a**) NS units tend to have higher mean firing rates than BS units. (**b**) By subsampling spikes from NS units with firing rates above the mean, a pseudopopulation was created with equivalent mean firing rate to BS units. (**B**) Summary of visual onset firing rate responses by cortical depth for the spike-equated NS and BS unit populations. (**a**) Scatter plot depicting visual onset responses for each unit at its estimated fractional depth. (**b**) Mean effect size (plus 99% confidence interval) across all recorded units (significant and non-significant visual onset responses included) by unit type and depth. No significant differences were observed between BS and spike-equated NS units. One-way ANOVA: ns p>0.05. (**c**) Histograms indicating percentages of units with significant excited and suppressed responses by unit type and depth.

Because stimulus offset responses have been observed in primary visual cortex (Liang et al., 2008), we also examined the possibility of significant firing rate changes following termination of the visual CMN stimulus in the current study. However, after FDR correction, we found that there were no units with significant visual offset responses. We therefore relied on visual onset responses throughout the remainder of the analyses to define visual responsive units.

### Visual-responsive neurons in A1 also typically respond to sound

The visual-responsive example unit shown in **Figure 2B** also had an apparent increase in firing rate aligned to sound onset, whereas the example in **Figure 2A** lacked any discernible change in firing rate during sound presentation. To quantify the proportions of unimodal and bimodal visual-responsive units, we examined intersections of significant firing rate changes evoked by unimodal visual and auditory stimuli. Because numerous prior studies reported differences between auditory transient spike rate changes at stimulus onset and sustained responses throughout the remainder of the stimulus, we separately analyzed firing rate responses averaged within the first 100 ms of the stimulus (onset) and from 200-ms to stimulus end (sustained). Example visual-responsive neurons with significant auditory onset and sustained responses are shown in **Figure 3A and B**, respectively. Consistent with previous studies, we found that onset responses were substantially stronger than sustained responses (**Figure 3C**) for both unit types (BS units: shallow: F = 8.25, p = 0.005, *η*^2^ = 0.053; middle: F = 32.68, p < 10^− 7^, *η*^2^ = 0.051; deep: F = 5.13, p = 0.025, *η*^2^ = 0.012; NS units: shallow: F = 33.15, p < 10^−6^, *η*^2^ = 0.274; middle: F = 40.98, p < 10^−8^, *η*^2^ = 0.170; deep: F = 26.17, p < 10^−5^, *η*^2^ = 0.168). As shown in **Figure 3D–E**, roughly half of all visual responsive units had either significant onset or sustained firing rate responses to sound. Considering onset and sustained responses together, well over two-thirds of visual-responsive neurons exhibited significant sound-evoked firing rate changes (**Figure 3F**).

**Figure 3.**
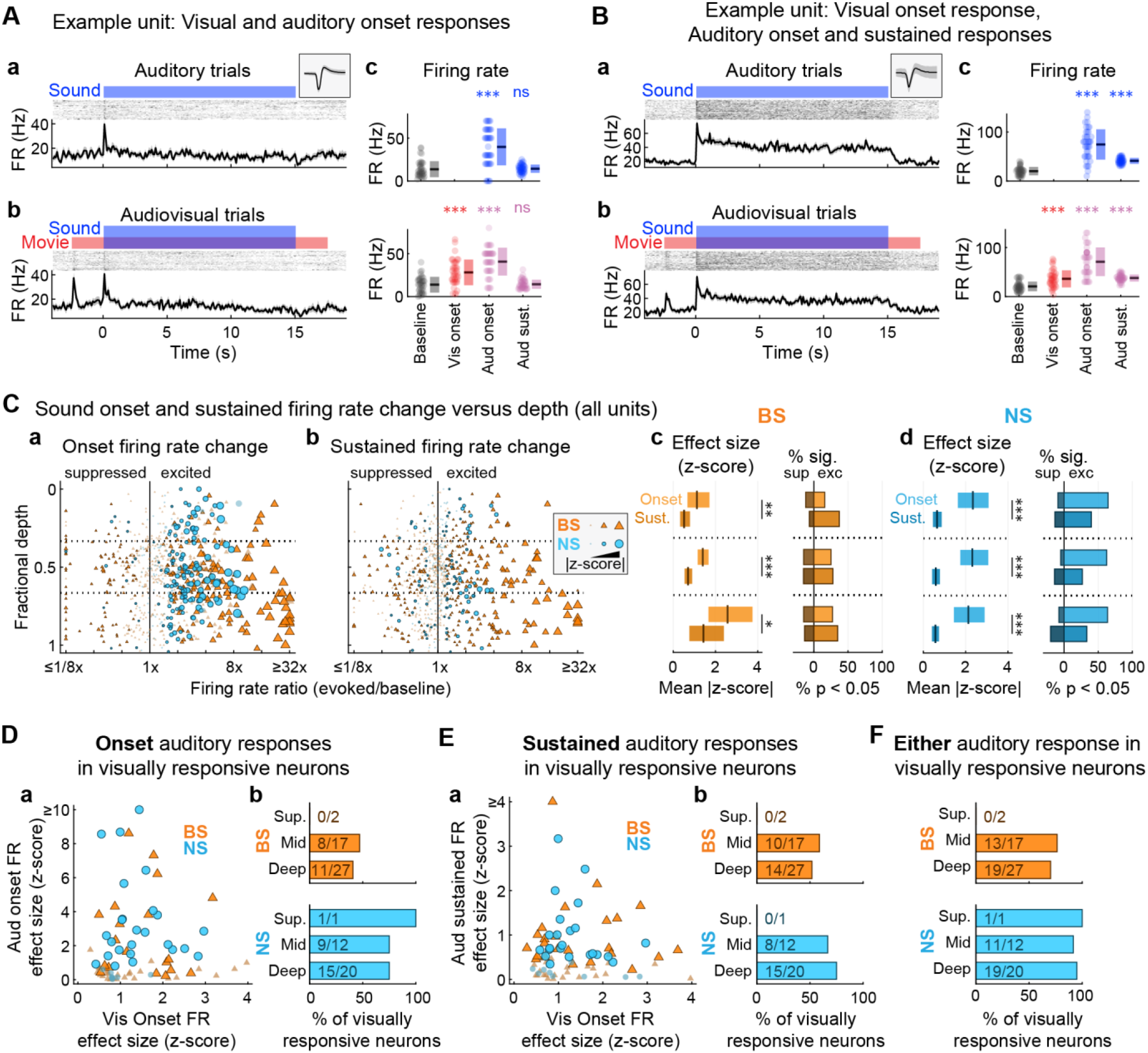
Visually responsive neurons in A1 also typically exhibit sound-evoked firing rate responses. (**A**) Example unit with visual and auditory onset evoked firing rate responses. (**a**) Auditory and (**b**) Audiovisual trials. Spiking responses quantified by peristimulus time histograms (lower) and binned-spike count matrices (upper) with red and blue bars indicating auditory and visual stimulus intervals, respectively (temporal binning: 100 ms). Inset in (a) shows the unit spike waveform (median ± MAD). (**c**) Window-averaged firing rate responses, with dots representing single trials and mean ± SD across trials indicated to the right. Wilcoxon signed-rank tests (paired): *p<0.05, **p<0.01, ***p<0.001, ns p>0.05. (**B**) Example unit with visual onset response as well as both auditory onset and sustained responses, with subplot organization as in (A). (**C**) Sound onset firing rate responses are stronger than sustained responses. (**a, b**) Summary of sound onset and sustained firing rate changes from baseline (including both visual responsive and non-responsive units) separated by unit type and cortical depth. (**c**) Comparison of onset and sustained responses for BS units. Left: mean effect size (plus 99% confidence interval) across all recorded units (significant and non-significant responses included). Effect sizes were significantly greater for onset responses (upper bars, lighter coloring) than sustained responses at each depth bin. One-way ANOVA: *p<0.05, **p<0.01, ***p<0.001, ns p>0.05. Right: Percentages of all recorded units with significant onset and sustained responses by unit type and depth. (**d**) Comparison of onset and sustained responses for NS units, with subplot organization as in (c). (**D**) Approximately half of visually responsive units have significant sound onset firing rate responses. (**a**) Scatter plot of visual and sound onset response effect sizes. Large markers with outlines reflect units with significant sound onset firing rate responses. (**b**) Bar plot showing the percentages of visually responsive units with significant sound onset firing rate responses. (**E**) Approximately half of visually responsive units have significant sound sustained firing rate responses. Subplot organization as in (D). (**F**) The majority of visually responsive units have significant sound-evoked firing rate responses including either onset or sustained responses. Bars represent the union of sound-responsive units E, b and D, b. Effect sizes were similarly greater for onset responses across depth bins.

In addition to onset and sustained responses, we observed transient firing rate changes at the offset of RDS segments in only a small minority of units (NS units: 15/212 [7.1%]; BS units: 27/589 [4.6%]). We thus focused on onset and sustained responses to define sound-evoked firing rate changes in the current report. Indeed, as in previous studies (Scholl et al., 2010), we found that the majority of these units also had significant onset and/or sustained responses (NS units: 12/15 [80.0%]; BS units: 14/27 [51.9%]).

The distinction between onset and sustained responses observed in the present study reinforces the conclusions of numerous previous studies suggesting spike rate changes driven by different stimulus phases (e.g., onset, offset, sustained) are at least partially dissociable and may contain different information about sounds (Lu et al., 2001; Wang et al., 2005; Malone et al., 2015). We pursued a related question as to whether sound-evoked responses reflecting spike rate and timing changes were similarly partially independent by examining intersections of sound-evoked firing rate responses (onset and sustained) and STRFs, which are sensitive to both spike rate and timing changes. As depicted by **Figure 4A, a–c**, STRFs are calculated by averaging the windowed stimulus segments preceding each spike. Thus, spectrotemporal tuning depends strictly upon temporal alignment between spike events and stimulus features. Importantly, such alignment may occur with or without changes from the spontaneous spike rate. We used a response reliability metric to determine whether observed STRF structure was statistically different from chance, as defined by STRFs calculated using time-reversed RDS segments (**Figure 4A, d–e**; Escabí et al., 2014).

**Figure 4.**
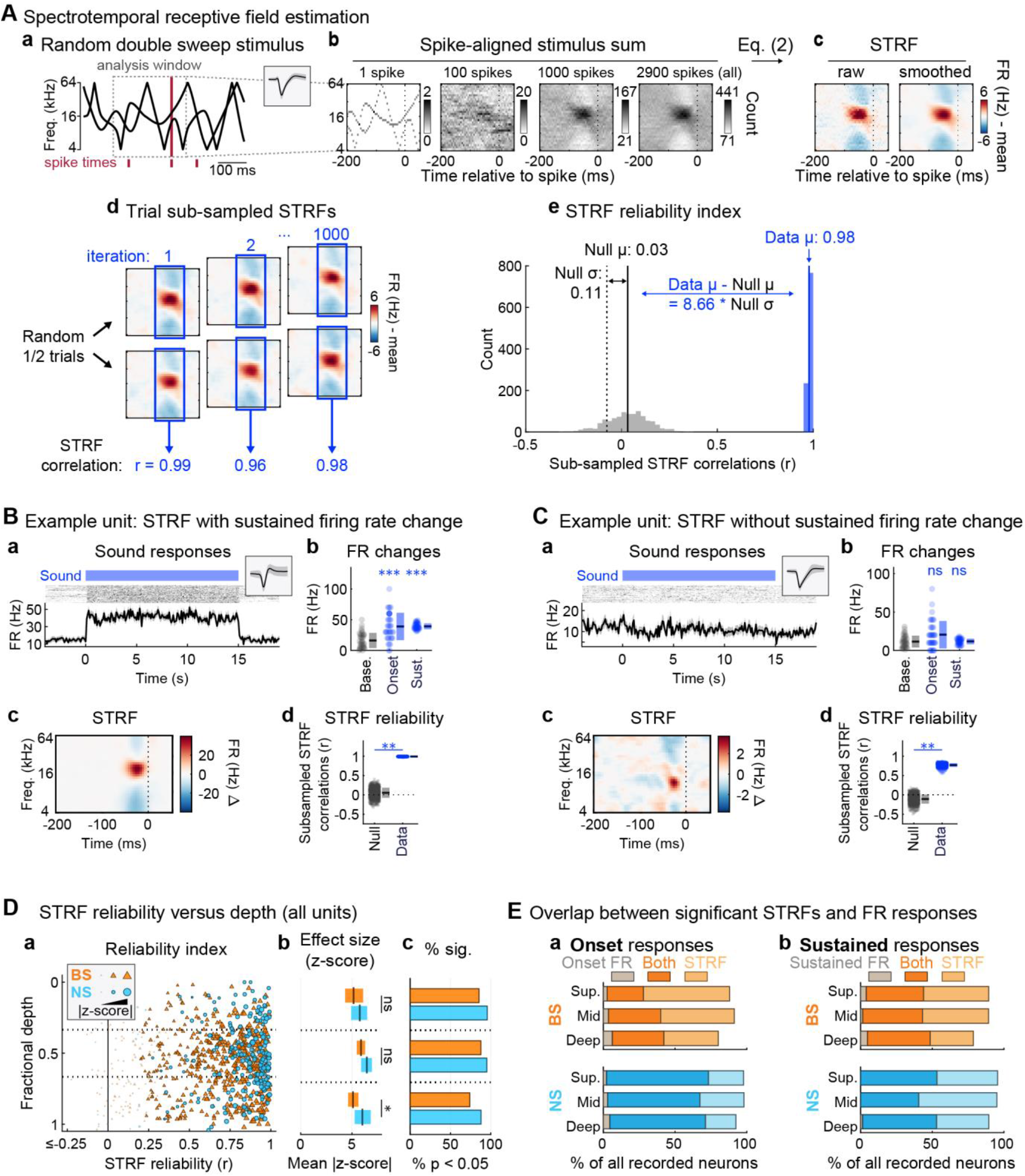
Auditory cortical neurons may encode spectrotemporal features with or without significant spike rate changes. (**A**) Spectrotemoral receptive field (STRF) estimation procedure. (**a**) For each spike, a segment of the RDS stimulus was stored using a window 200 ms before and 50 ms after the spike time. (**b**) RDS segments aligned to each spike were added to a cumulative sum. Structure in the time-frequency bins typically only emerges after several hundred spikes or more. (**c**) Transforming the spike-aligned stimulus sum according to Eq. (2) yielded an STRF estimate expressed in firing rate (Hz) deviations from the mean driven rate. Red and blue regions indicate stimulus energy at time-frequency bins associated with increases and decreases in firing rate, respectively. (**d**) A subsampling procedure was used to determine the statistical significance of time-frequency bin structure in the STRFs. The correlation coefficient between STRFs calculated from random trial halves (without replacement) was calculated across 1000 iterations. (**e**) The STRF reliability index was defined as the mean of the subsampled correlation coefficient distribution. A null distribution was obtained from STRFs calculated using time-reversed stimulus RDS segments, which breaks the temporal relationship between spikes and stimulus features but preserves spike count and timing statistics. A p-value was obtained by dividing the number of null STRF correlations exceeding the reliability index (data) by the number of iterations and multiplying by two for two-tailed significance. Effect size reflected the absolute difference between null and data means, divided by the null standard deviation. (**B**) Example unit with significant STRF reliability as well as onset and sustained firing rate responses. (**a**) Spiking responses quantified by peristimulus time histograms (lower) and binned-spike count matrices (upper) with blue bar indicating auditory stimulus interval (temporal binning: 100ms). Inset shows the unit spike waveform (median ± MAD). (**b**) Summary of sound onset and sustained firing rate responses compared to baseline. Each dot represents a single trial, with mean ± SD across trials indicated to the right. Wilcoxon signed-rank tests (paired): *p<0.05, **p<0.01, ***p<0.001, ns p>0.05. (**c**) STRF as calculated in (A, a–c). (**d**) STRF reliability as calculated in (A, d–e). Each dot represents the correlation between STRFs for a single subsample iteration, with mean ± SD across trials indicated to the right. Subsampling test: *p<0.05, **p<0.01, ***p<0.001, ns p>0.05. (**C**) Example unit significant STRF reliability but non-significant onset and sustained firing rate responses. Subplot organization as in (B). (**D**) Summary of STRF reliability by unit type and cortical depth. (**a**) STRF reliability for each unit at its estimated cortical depth. Marker sizes are scaled by effect size, with outlined markers indicating units with significant STRF reliability (p < 0.05, Benjamini– Hochberg FDR correction). (**b**) Mean effect size (plus 99% confidence interval) across all recorded units (units with significant and non-significant reliability included) by unit type and depth. A small difference between unit types was observed in the deepest cortical bin. One-way ANOVA: *p<0.05, **p<0.01, ***p<0.001, ns p>0.05. (**c**) Histograms indicating percentages of all recorded units with significant STRF reliability. (**E**) Significant STRF reliability may occur with or without significant firing rate changes. (**a**) Intersections of significant onset firing rate responses alone, significant STRF reliability alone, or both. (**b**) Intersections of significant sustained firing rate responses alone, significant STRF reliability alone, or both.

An example unit with significant STRF reliability, which also had clear firing rate changes from baseline, is shown in **Figure 4B**. By contrast, **Figure 4C** shows an example unit with significant STRF reliability but neither significant onset nor sustained firing rate changes. Significantly reliable STRFs were observed in the majority of both NS and BS units (**Figure 4D**). Notably, significant changes in firing rate were not observed in many of the units with significant STRFs (**Figure 4E**). For BS units, significant STRFs without onset responses were slightly more common than both STRF and onset responses together, whereas the reverse was true for NS units (**Figure 4E, a**). For both unit types, sustained responses were observed in roughly half of units with significant STRFs (**Figure 4E, b**). Significant firing rate changes (onset or sustained) without STRFs were rare. Because sustained responses and STRFs were calculated from the same analysis windows, spikes, and stimulus distributions, our results underscore the important distinction between changes in spike rate and timing in response to sound.

We quantified how many visual-responsive neurons were also responsive to sound features as revealed by STRF calculation, e.g., as shown by the visual-responsive example unit with significant spectrotemporal tuning in **Figure 5A**. Group data indicated that visual-responsive neurons with significant STRFs were the rule rather than the exception, with significant STRFs observed in all but a handful of BS units (**Figure 5B**). Extending the definition of ‘sound responsive’ to include STRFs as well as onset or sustained firing rate responses indicated that, with just a few exceptions, visual-responsive neurons in A1 also respond to sound (**Figure 5C**).

**Figure 5.**
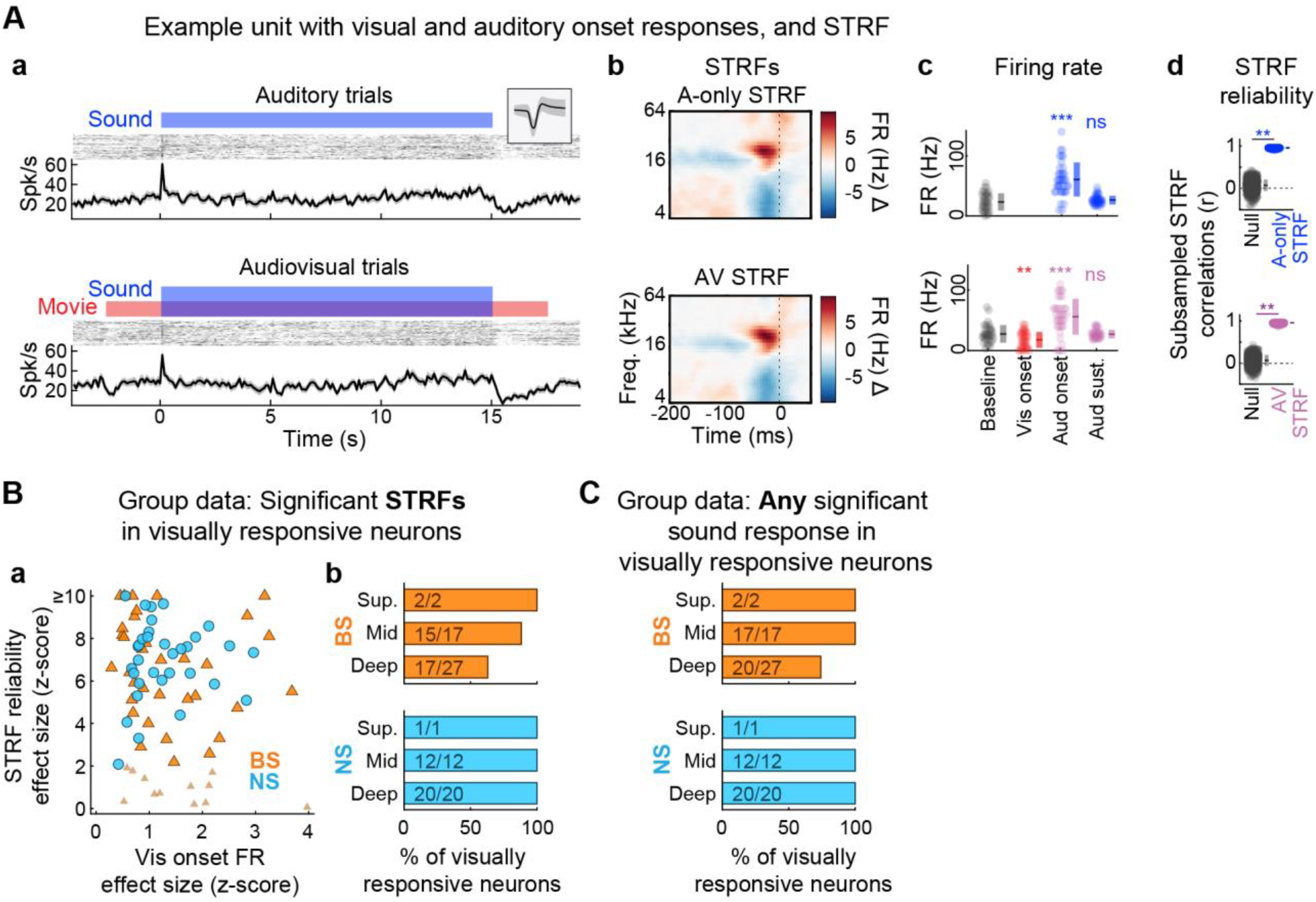
Visually responsive A1neurons are typically tuned for spectrotemporal features. (**A**) Example unit with both visual and auditory onset firing rate responses as well as significant STRF reliability. (**a**) Spiking responses quantified by peristimulus time histograms (lower) and binned-spike count matrices (upper) with blue and red bars indicating auditory and visual stimulus intervals, respectively (temporal binning: 100ms). Inset shows the unit spike waveform (median ± MAD). (**b**) STRFs for each condition. (**c**) Summary of firing rate responses compared to baseline. Each dot represents a single trial, with mean ± SD across trials indicated to the right. Wilcoxon signed-rank tests (paired): *p<0.05, **p<0.01, ***p<0.001, ns p>0.05. (**d**) STRF reliability for each condition. Each dot represents the correlation between STRFs for a single subsample iteration, with mean ± SD across trials indicated to the right. Subsampling test: *p<0.05, **p<0.01, ***p<0.001, ns p>0.05. (**B**) The majority of visually responsive units have significant STRF reliability. (**a**) Scatter plot showing effect size for visual onset and STRF reliability. The larger, outlined markers indicate units with significant STRF reliability. (**b**) Bar plots showing percentages of visually responsive units with significant STRF reliability. (**C**) The majority of visually responsive units also respond to sound as defined by either significant firing rate responses (onset or sustained) or STRF reliability. Bars represent the union of units with significant STRF reliability and units with significant firing rate responses (onset or sustained).

The preceding results indicated that if a neuron was responsive to visual stimuli, it was likely also responsive to sound. We addressed a corollary question of whether the presence or absence of an auditory response predicted whether a unit was responsive to visual stimuli. Chi-squared tests were used to compare percentages of visual-responsive units within unit subpopulations separated by significant and non-significant auditory responses. As depicted in **Figure 6**, visual responsiveness was not strongly predicted by the presence or absence of any auditory response type for either unit subpopulation. Small but statistically significant effects were observed for NS units in the deepest bin for sustained spike rate changes (**Figure 6B**; Χ^2^ = 4.63, p = 0.031) and STRFs (**Figure 6C**; Χ^2^ = 4.28, p = 0.039). A similar trend was observed for sustained responses in the middle depth bin (**Figure 6B**; Χ^2^ = 3.84, p = 0.050). These results suggested that the absence of either sound response was associated with decreased probability of a visual response, recapitulating analyses above suggesting responses to the two modalities tended to occur together. Differences for all other depth bins, and across depth bins for BS units were non-significant (all Χ^2^ < 1.90, all p-values > 0.16). Thus, with minor exceptions, visual responses were approximately evenly distributed among sound response types, leaving cortical depth as the most meaningful predictor of visual responsiveness.

**Figure 6.**
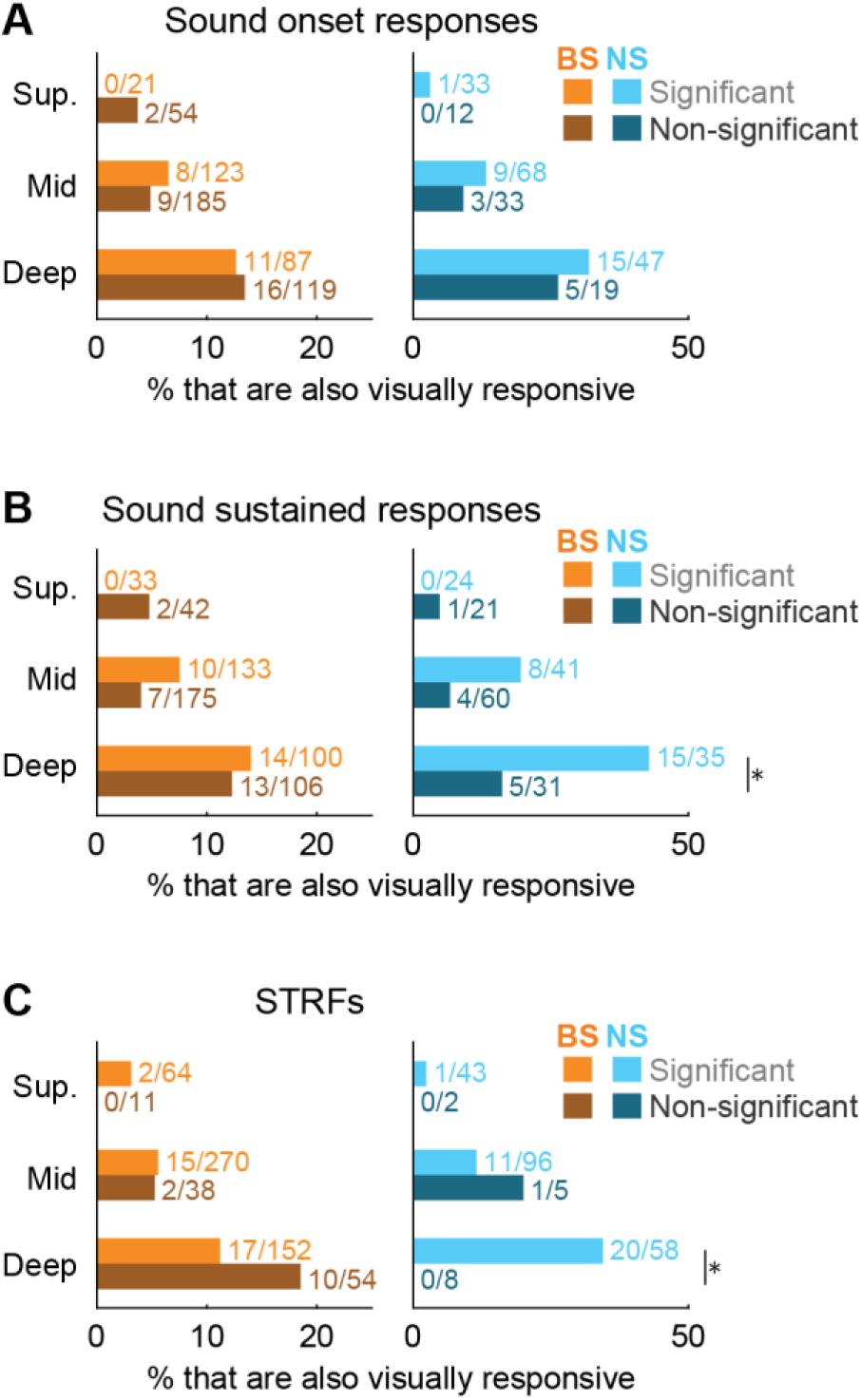
Sound response archetypes are not strongly predictive of visual responsiveness. Each subplot shows the proportion of BS and NS units (left and right) separated by cortical depth with and without significant sound responses (light and dark bars) as defined by (**A**) onset firing rate responses, (**B**) sustained firing rate responses, and (**C**) STRFs. NS units in the deepest cortical bin with significant sustained sound responses and STRFs were more likely to have significant visual responses (Chi-square tests: *p<0.05). The presence or absence of any other sound response at any other depth bin did not predict visual responsiveness.

### Visual context differentially modulates sound onset and sustained firing rate responses

The foregoing results documented unimodal visual responses in a subpopulation of A1 neurons, and further that most of these units were bimodal, responding to both visual and auditory stimuli. We investigated further the extent of audiovisual integration in A1 by examining the frequency with which sound onset and sustained firing rate responses were themselves modulated by the presence or absence of the visual stimulus. **Figure 7A** shows an example neuron for which both sound onset and sustained responses significantly increased on audiovisual trials. For the example unit in **Figure 7B**, the sustained response decreased significantly on audiovisual trials, but the onset was unaffected. The latter outcome was typical of units with visual-modulated responses to sound. Indeed, the example neuron in **Figure 7A** was the only unit in the entire population for which sound onset response changed significantly with visual stimulation (**Figure 7C, a**). By comparison, sustained sound firing rate responses were significantly modulated by visual context in 7.1% of NS units and 5.1% of BS units (**Figure 7C, b**). ANOVA confirmed visual modulation effect sizes were larger for sustained than onset responses for the middle and deep cortical depth bins for both unit types (BS units: shallow: F = 0.16, p = 0.687, *η*^2^ = 0.001; middle: F = 8.62, p = 0.003, *η*^2^ = 0.014; deep: F = 9.79, p = 0.002, *η*^2^ = 0.024; NS units: shallow: F = 0.10, p = 0.756, *η*^2^ = 0.001; middle: F = 6.60, p = 0.011, *η*^2^ = 0.032; deep: F = 9.46, p = 0.003, *η*^2^ = 0.068). Notably, these outcomes were exactly opposite of responses driven by sound alone, in which case onsets were substantially stronger than sustained firing rate changes (**Figure 3**).

**Figure 7.**
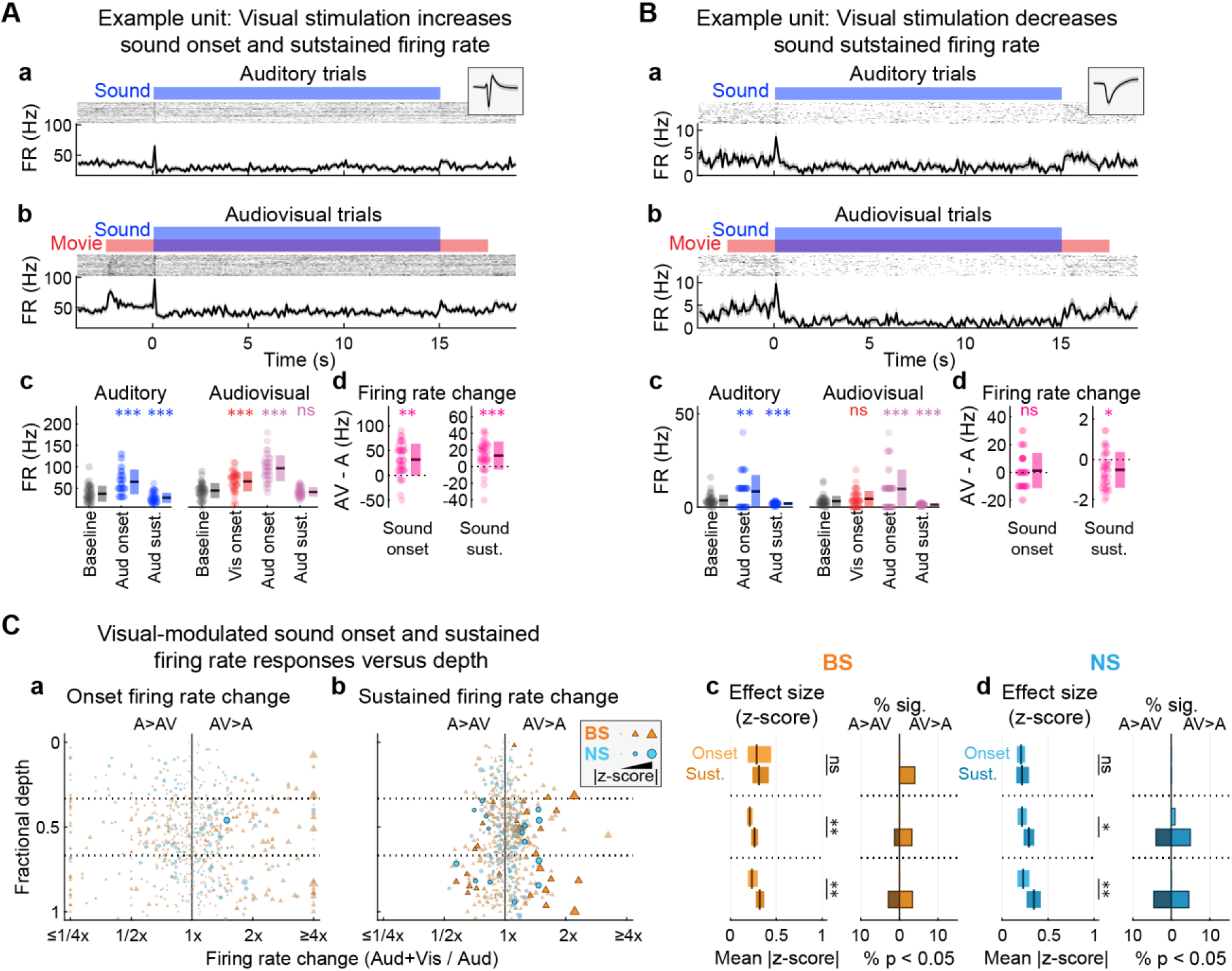
Visual context differentially modulates sound onset and sustained firing rate responses. (**A**) Example unit for which visual stimulation significantly increased onset and sustained firing rate responses to sound. (**a**) Auditory and (**b**) Audiovisual trials. Spiking responses quantified by peristimulus time histograms (lower) and binned-spike count matrices (upper) with red and blue bars indicating auditory and visual stimulus intervals, respectively (temporal binning: 100 ms). Inset in (a) shows the unit spike waveform (median ± MAD). (**c**) Window-averaged firing rate responses, with dots representing single trials and mean ± SD across trials indicated to the right. Wilcoxon signed-rank tests (paired): *p<0.05, **p<0.01, ***p<0.001, ns p>0.05. (**d**) Difference in averaged firing rate responses between conditions (Audiovisual - Auditory trials), with dots representing single trials and mean ± SD across trials indicated to the right. Wilcoxon signed-rank tests (paired): *p<0.05, **p<0.01, ***p<0.001, ns p>0.05. (**B**) Example unit for which visual stimulation significantly increased sustained but not onset firing rate responses to sound. Subplot organization as in (A). (**C**) Visual modulation effects are stronger for sound sustained firing rate responses than onset responses. (**a, b**) Summary of visual-modulated sound onset and sustained firing rate changes (Audiovisual divided by Auditory) separated by unit type and cortical depth for all units. Scatter plots depict firing rate changes between conditions for each unit by cortical depth. Firing rate ratio values above and below 1 (x-axis) indicate increases and decreases in firing on Audiovisual trials relative to Auditory trials, respectively. Outlined markers indicate statistically significant responses. Marker sizes are scaled by effect size (absolute difference between Audiovisual and Auditory means, divided by Auditory SD). (**c**) Comparison of visual-modulated onset and sustained responses for BS units. Left: mean effect size (plus 99% confidence interval) across all recorded units (significant and non-significant responses included). Effect sizes were significantly greater for sustained responses (lower bars, darker coloring) than onset responses at the middle and deep bins. One-way ANOVA: *p<0.05, **p<0.01, ***p<0.001, ns p>0.05. Right: Percentages of all recorded units with significant visual-modulated onset and sustained responses. (**d**) Comparison of onset and sustained responses for NS units, with subplot organization as in (c). Effect sizes were similarly greater for sustained responses (lower bars, darker coloring) at the middle and deep bins. See Extended Data Figure 7-1 for results separated by units for which sound responses reflected increases and decreases from baseline.

Consistent with the concentration of unimodal visual responses in the deepest cortical bin (**Figure 2**), the majority of units with visual-modulated sustained responses to sound were observed in either the middle or deep cortical bin, with the strongest mean effect size in the deepest bin (**Figure 7C, c–d**). ANOVA confirmed small but significant effects of cortical depth on visual modulated sustained sound responses for both unit types (BS units: F = 3.22, p = 0.041, *η*^2^ = 0.011; NS units: F = 4.69, p = 0.010, *η*^2^ = 0.043). Visual modulated onset responses were not significantly dependent upon depth (BS units: F = 2.35, p = 0.096, *η*^2^ = 0.008; NS units: F = 0.25, p = 0.780, *η*^2^ = 0.002). All differences between unit type were similarly non-significant (all F-ratios < 1.5, p-values > 0.23), with the exception of borderline effect suggesting stronger sustained response modulation for BS units in the shallowest bin (F = 3.91, p = 0.051, *η*^2^ = 0.032).

Sound-evoked sustained firing rates were suppressed below baseline for the example units in **Figure 7A–B**. However, the effect of visual stimulation was opposite for each neuron, elevating firing near the spontaneous rate for the example in **Figure 7A** and further decreasing the suppressed response for the example in **Figure 7B**. These examples raised the possibility that effects of visual stimulation on sound-evoked firing rate responses might differ depending on whether responses to sound alone reflected an increase or decrease in spike rate relative to baseline (spontaneous). We thus expanded the analyses above by dividing onset and sustained responses into subgroups for which the sound-evoked response was greater or less than spontaneous. Distributions of sound-evoked firing rate responses divided by baseline rates are shown in **Extended Data Figure 7-1A–B**, indicating both increases and decreases in firing rate relative to baseline were common for both onset and sustained responses in both unit subpopulations. Designations of ‘excited’ and ‘suppressed’ units were inclusive of both significant and non-significant increases and decreases in firing rate, respectively, to accommodate potential cases in which sound-evoked responses only reached significance in one or the other condition (e.g., sustained response in **Figure 7A**). By separating visual-modulatory influences into excited and suppressed subgroups, the analysis included four possible forms of modulation: increases and decreases in excited and suppressed responses.

**Extended Data Figure 7-1.**
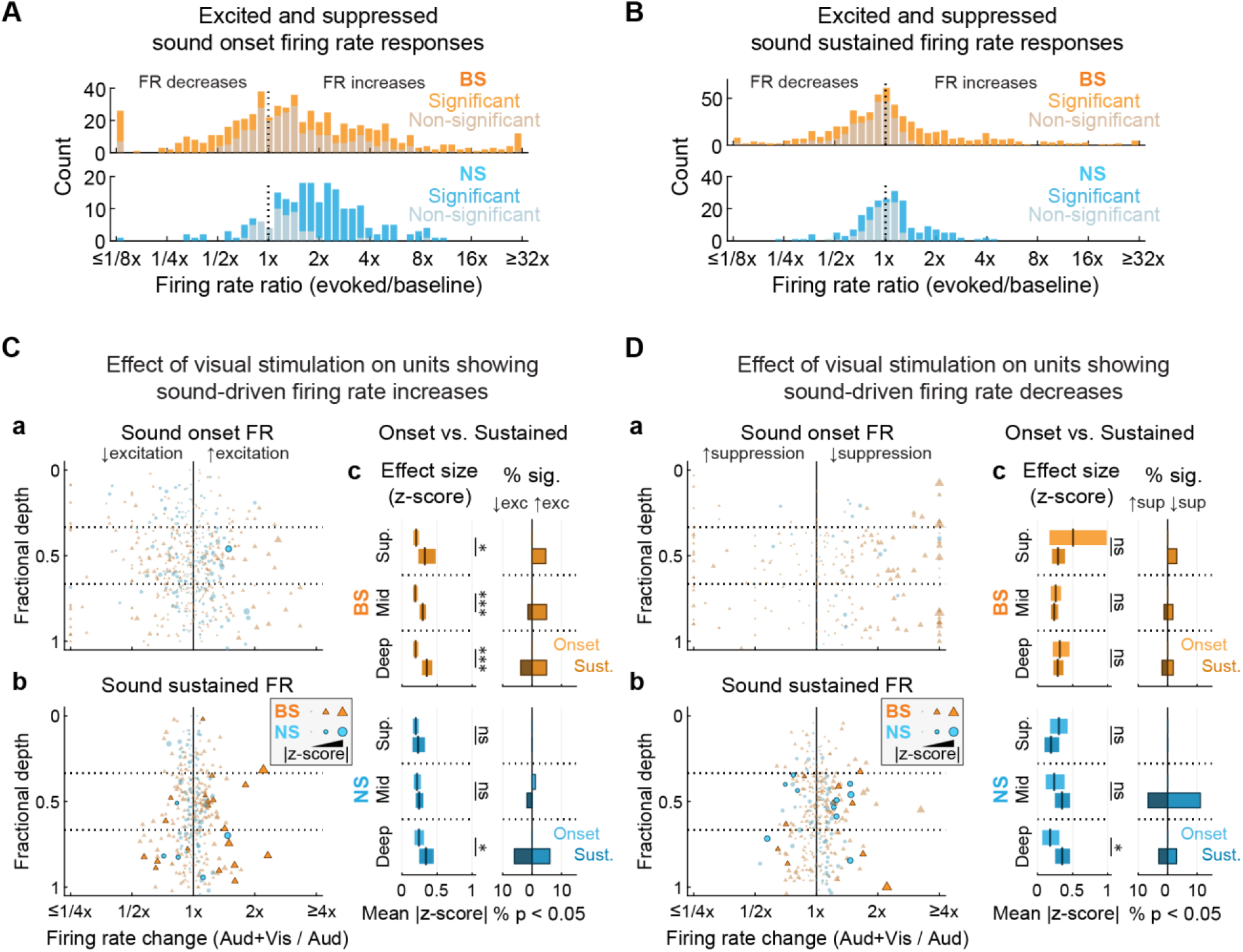
Visual modulation of excited and suppressed sound onset and sustained firing rate responses. (**A**) Sound may evoke increases or decreases in onset firing rate. Histograms show the numbers of units (BS, top; NS, bottom) with increases and decreases in firing rate relative to baseline, with significant (p<0.05) and non-significant (p>=0.05) responses indicated by dark and light bars, respectively. (**B**) Sound may evoke increases or decreases in sustained firing rate. Histograms as in (A). (**C**) Summary of visual stimulation effects on onset and sustained firing rate responses that were excited relative to baseline. (**a, b**) Summary of visual-modulated sound onset and sustained firing rate changes (Audiovisual divided by Auditory) separated by unit type and cortical depth for all units with excited responses. Scatter plots depict firing rate changes between conditions for each unit by cortical depth. Firing rate ratio values above and below 1 (x-axis) indicate increases and decreases in excitation on Audiovisual trials relative to Auditory trials, respectively. Outlined markers indicate statistically significant responses. Marker sizes are scaled by effect size (absolute difference between Audiovisual and Auditory means, divided by Auditory SD). (**c**) Comparison of visual-modulated onset and sustained responses for BS (top) and NS units (bottom). Left: mean effect size (plus 99% confidence interval) across units with excited sound responses. Visual modulation effect sizes were significantly greater for sustained responses (lower bars, darker coloring) than onset responses across depth bins for BS units, and at the deepest bin for NS units. One-way ANOVA: *p<0.05, **p<0.01, ***p<0.001, ns p>0.05. Right: Percentages of all recorded units with significant visual-modulated onset and sustained responses. (**D**) Summary of visual stimulation effects on onset and sustained firing rate responses that were suppressed relative to baseline. Subplot organization as in (C).

Outcomes of this analysis were generally consistent with the pooled results reported above. For BS units with excited responses (**Extended Data Figure 7-1C, c**), visual modulation effects were stronger for sustained than onset responses at all depth bins (shallow: F = 5.99, p = 0.016, *η*^2^ = 0.062; middle: F = 27.20, p < 10^−6^, *η*^2^ = 0.074; deep: F = 29.82, p < 10^−6^, *η*^2^ = 0.115). For BS units with suppressed responses (**Extended Data Figure 7-1D, c**), differences in visual modulation between sustained and onset responses were non-significant at all depth bins (all F-ratios < 2.15, p-values > 0.14). These non-significant outcomes may reflect floor effects limiting suppression of the already relatively low spontaneous rates in BS units. For NS units with both excited and suppressed responses, visual modulation effects were stronger for sustained than onset responses at the deepest bin only (excited: F = 4.59, p = 0.035, *η*^2^ = 0.051; suppressed: F = 4.79, p = 0.034, *η*^2^ = 0.100; all other F-ratios < 2.0, p-values > 0.16).

For the example neuron in **Figure 7A**, but not the example in **Figure 7B**, visual modulated responses to sound coincided with a significant response to visual stimulation alone. We examined the consistency of these outcomes by calculating intersections between unimodal visual responses and visual-modulated sustained responses to sound. **Figure 8A** shows an example unit for which a sustained, excitatory response to sound was further elevated by visual stimulation even though the response to visual stimulation alone was not significant. This was true for the majority of units with visual-modulated sustained sound responses, as seen in **Figure 8B**, which did not respond outright to unimodal visual stimuli. This finding suggests only partial overlap between visual responsive and visual modulated unit subpopulations.

**Figure 8.**
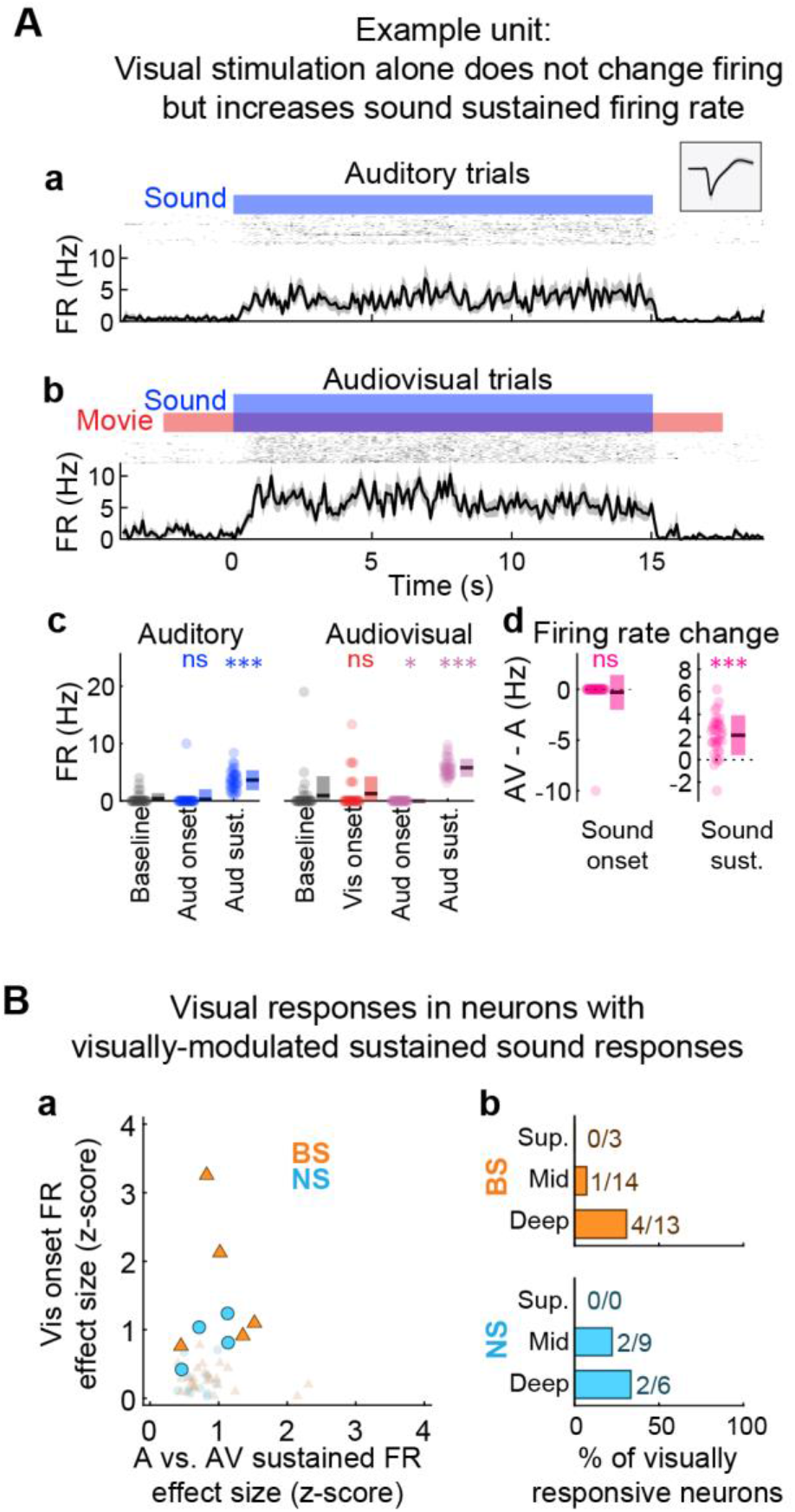
Sound-evoked firing rate responses may be modulated by visual inputs even without responses to visual stimulation alone. (**A**) Example unit with non-significant visual onset response but significant visual-modulated increases in sustained firing rate responses to sound. (**a**) Auditory and (**b**) Audiovisual trials. Spiking responses quantified by peristimulus time histograms (lower) and binned-spike count matrices (upper) with red and blue bars indicating auditory and visual stimulus intervals, respectively (temporal binning: 100 ms). Inset in (a) shows the unit spike waveform (median ± MAD). (**c**) Window-averaged firing rate responses, with dots representing single trials and mean ± SD across trials indicated to the right. Wilcoxon signed-rank tests (paired): *p<0.05, **p<0.01, ***p<0.001, ns p>0.05. (**d**) Difference in averaged firing rate responses between conditions (Audiovisual - Auditory trials), with dots representing single trials and mean ± SD across trials indicated to the right. Wilcoxon signed-rank tests (paired): *p<0.05, **p<0.01, ***p<0.001, ns p>0.05. (**B**) Many units with significant visual-modulated sustained sound firing rate responses are not responsive to visual stimulation alone. (**a**) Scatter plot of effect sizes for visual onset responses and visual-modulated sustained sound responses. Large markers with outlines reflect units with significant visual onset firing rate responses. (**b**) Bar plot showing the percentages of units with significant visual-modulated sustained sound responses that are also visual responsive.

### Visual stimulation may modify STRFs independently of firing rate changes

As reported above, many units for which sound-evoked firing rate responses were significantly modulated by visual stimulation were not significantly responsive to visual stimulation alone. This suggested that STRFs might likely also be modulated by the presence of visual stimuli, possibly for units without responses to unimodal visual stimuli. We therefore examined visual influences on spectrotemporal encoding by calculating difference STRFs, referred to hereafter as ΔSTRFs, reflecting the STRF in the auditory condition subtracted from the STRF in the audiovisual condition. The significance of ΔSTRFs was determined by iteratively subsampling STRFs from each condition and calculating the difference, then comparing the ΔSTRF correlation distribution to an equivalent null distribution (**Figure 9A, a–b**). An example neuron with significant ΔSTRF reliability is shown in **Figure 9B**. In total, 7.1% of NS units and 6.3% of BS units had STRFs that were significantly modulated by visual stimulation (**Figure 9C**). There was no evidence that ΔSTRF reliability depended significantly on cortical depth or unit type (all F-ratios < 2.6, all p-values > 0.11). As with sustained firing rate responses to sound, visual modulation of STRFs most often occurred without significant responses to visual stimulation alone (**Figure 9D**).

**Figure 9.**
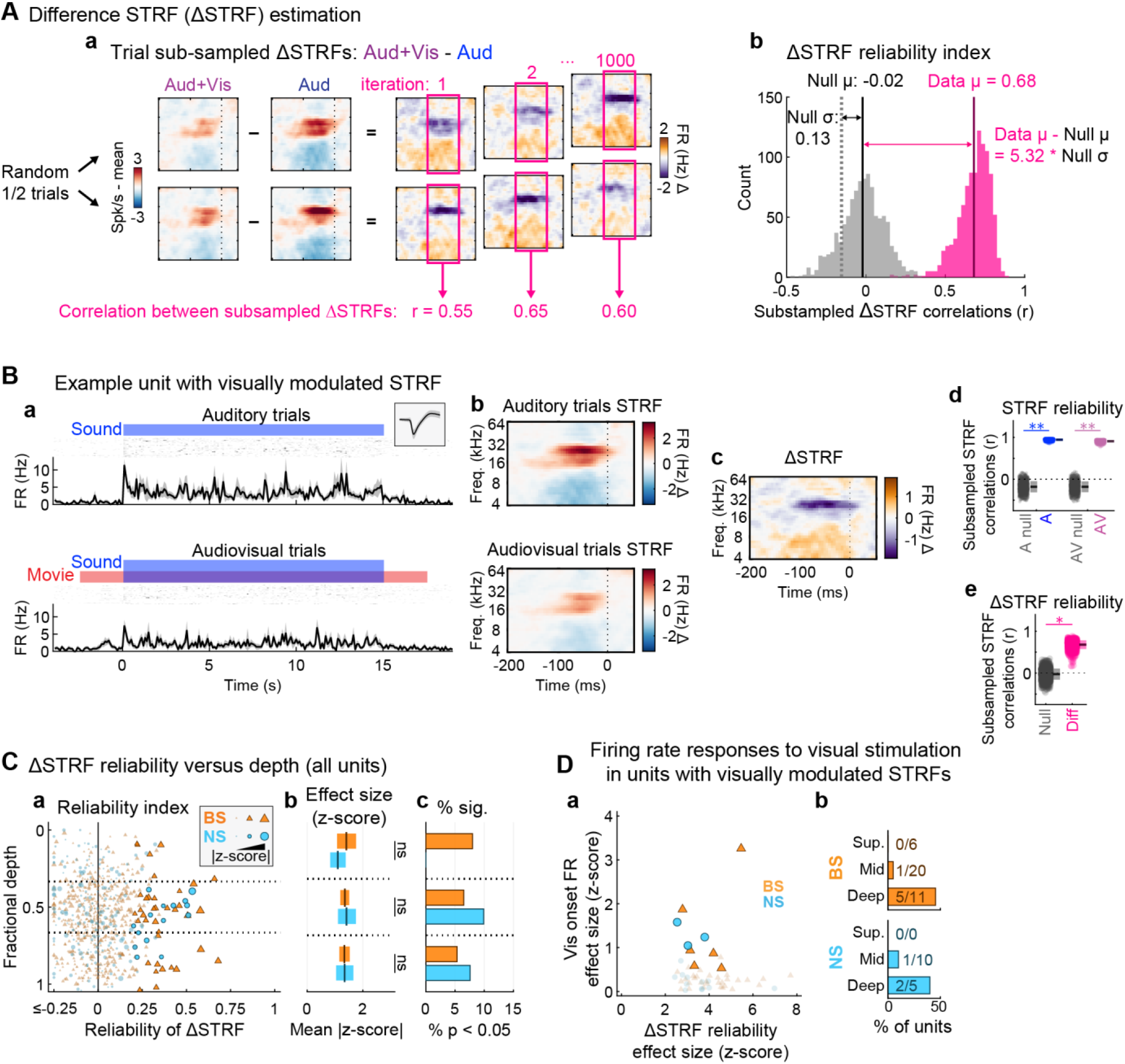
Visual context may modify spectrotemporal receptive fields even in units without significant responses to visual stimulation alone. (**A**) Procedure for estimating differences in STRFs between conditions (ΔSTRFs). (**a**) ΔSTRFs were estimated by subtracting the Audiovisual STRF from the Auditory alone STRF. A subsampling test was used to determine the statistical significance of time-frequency bin structure in the ΔSTRFs. The correlation coefficient between ΔSTRFs calculated from random trial halves (without replacement) was calculated across 1000 iterations. (**b**) ΔSTRF reliability was defined as the mean of the subsampled correlation coefficient distribution. A null ΔSTRF distribution was obtained from ΔSTRFs calculated using time-reversed stimulus RDS segments. A p-value was obtained by dividing the number of null STRF correlations exceeding the reliability index (data) by the number of iterations and multiplying by two for two-tailed significance. Effect size reflected the absolute difference between null and data means, divided by the null standard deviation. (**B**) Example unit with significant ΔSTRF reliability. (**a**) Spiking responses quantified by peristimulus time histograms (lower) and binned-spike count matrices (upper) with blue and red bars indicating auditory and visual stimulus intervals, respectively (temporal binning: 100ms). Inset shows the unit spike waveform (median ± MAD). (**b**) STRFs for each condition (**c**) ΔSTRF (**d**) STRF reliability for each condition. (**e**) ΔSTRF reliability. For (d) and (e), each dot represents the correlation between STRFs or ΔSTRFs for a single subsample iteration, with mean ± SD across trials indicated to the right. Subsampling test: *p<0.05, **p<0.01, ***p<0.001, ns p>0.05. (**C**) Summary of ΔSTRF reliability by unit type and cortical depth. (**a**) ΔSTRF reliability for each unit at its estimated cortical depth. Marker sizes are scaled by effect size, with outlined markers indicating units with significant ΔSTRF reliability (p < 0.05, Benjamini–Hochberg FDR correction). (**b**) Mean effect size (plus 99% confidence interval) across all recorded units (units with significant and non-significant reliability included) by unit type and depth. No differences between unit types were observed. One-way ANOVA: ns p>0.05. (**c**) Histograms indicating percentages of all recorded units with significant ΔSTRF reliability. (**D**) Significant ΔSTRF reliability often occurs without significant visual onset firing rate changes. (**a**) Scatter plot of visual onset and ΔSTRF reliability response effect sizes. Large markers with outlines reflect units with significant visual onset firing rate responses. (**b**) Bar plot showing the percentages of units with significant ΔSTRF reliability that are also visually responsive.

Two outcomes reported above further raised the possibility that STRFs may be modulated by visual stimulation independently of firing rate changes. First, significant STRFs were frequently obtained on auditory trials even in the absence of time-averaged firing rate changes (**Figure 4**). Second, sound-evoked firing rate changes (onset vs. sustained) were themselves independently modulated by visual context (**Figure 7**). To address this possibility, we examined overlap between units with significant ΔSTRF reliability and visual-modulated sustained firing rate responses. For the example unit in **Figure 10A**, the STRF was significantly modulated by visual context, even though the sustained firing rate response did not significantly change from baseline, let alone between conditions. Group data presented in **Figure 10B** revealed that similar independent modulation of sustained firing rate responses and STRFs occurred in the majority of units sensitive to visual context.

**Figure 10.**
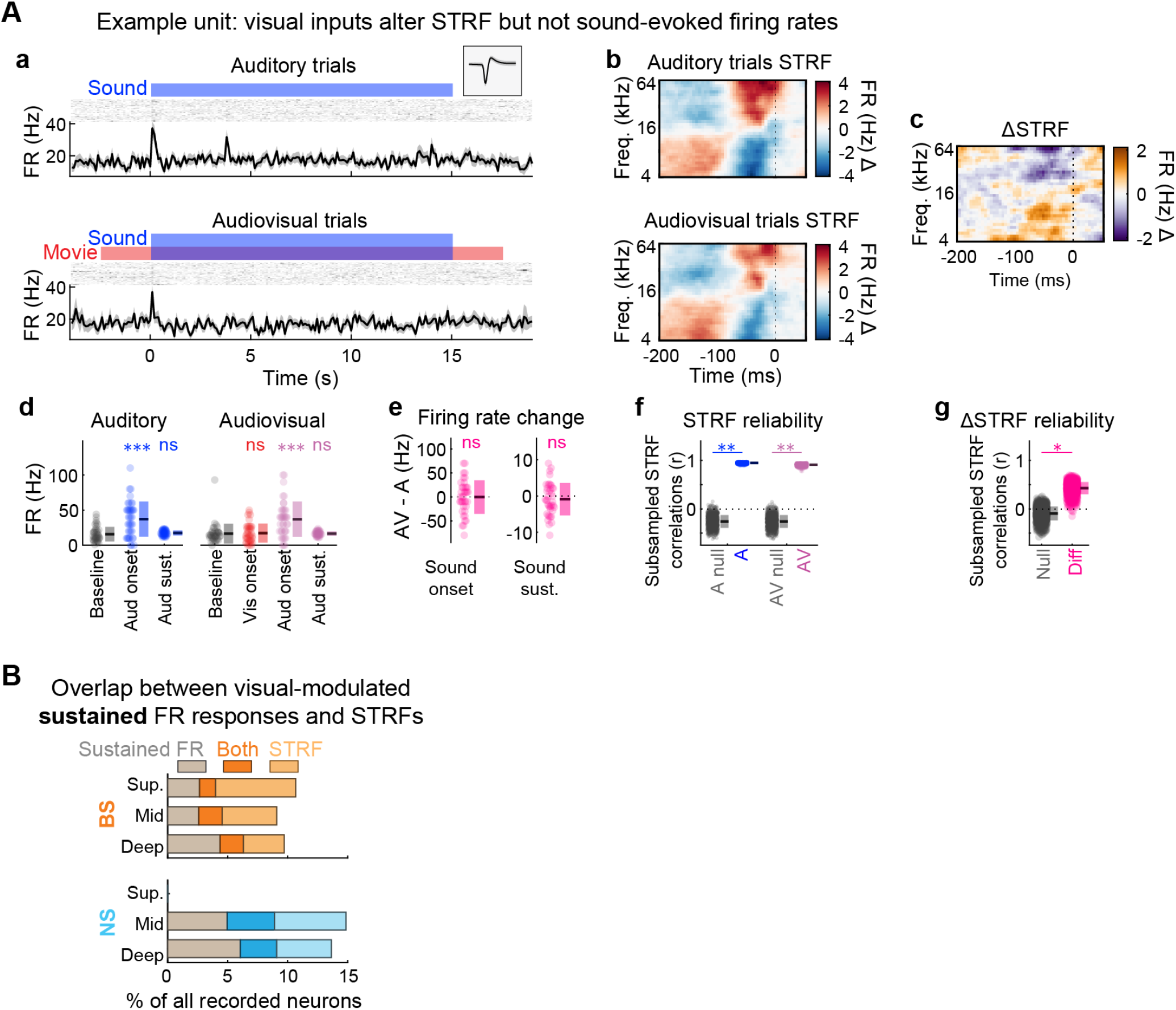
Visual stimulation may modify spectrotemporal receptive fields independently of averaged firing rates. (**A**) Example unit with significant ΔSTRFs reliability but non-significant visual-modulated sustained firing rate. (**a**) Spiking responses quantified by peristimulus time histograms (lower) and binned-spike count matrices (upper) for each condition with blue and red bars indicating auditory and visual stimulus intervals, respectively (temporal binning: 100ms). Inset shows the unit spike waveform (median ± MAD). (**b**) STRFs for each condition (**c**) ΔSTRF (**d**) Window-averaged firing rate responses, with dots representing single trials and mean ± SD across trials indicated to the right. Wilcoxon signed-rank tests (paired): *p<0.05, **p<0.01, ***p<0.001, ns p>0.05. (**e**) Difference in averaged firing rate responses between conditions (Audiovisual - Auditory trials), with dots representing single trials and mean ± SD across trials indicated to the right. Wilcoxon signed-rank tests (paired): *p<0.05, **p<0.01, ***p<0.001, ns p>0.05. (**f**) STRF reliability for each condition. (**g**) ΔSTRF reliability. For (f) and (g), each dot represents the correlation between STRFs or ΔSTRFs for a single subsample iteration, with mean ± SD across trials indicated to the right. Subsampling test: *p<0.05, **p<0.01, ***p<0.001, ns p>0.05. (**B**) Significant ΔSTRF reliability may occur with or without significant visual modulation of sustained sound-evoked firing rate changes. Bar plots indicate intersections of significant visual modulated sustained firing rate responses alone, significant ΔSTRF reliability alone, or both.

### Visual context may alter STRF gain but preserves spectrotemporal tuning

In the previous sections, we found that STRFs are sensitive to visual context for some units as indicated by the reliability of ΔSTRFs. This metric confirmed that the STRFs differed significantly between conditions but did not reveal what was different about the STRFs, qualitatively or quantitatively. We therefore examined the degree to which differences between conditions reflected gain (quantitative) or tuning changes (qualitative) by calculating slope and correlation coefficient parameters for time-frequency bin values between conditions. For this analysis, we included only STRFs for which the reliability index in the auditory condition was >0.5 to ensure reliable baseline spectrotemporal tuning prior to estimating differences between conditions. Example units with relatively strong and weak correlations between STRFs in the auditory and audiovisual conditions are shown in **Figure 11A, a and b**, respectively. As indicated by **Figure 11A, c**, STRFs were usually highly correlated between conditions (NS median: r = 0.95; BS median: r = 0.90), implying largely preserved spectrotemporal tuning between conditions. Absolute correlations between auditory STRFs and ΔSTRFs were similarly high (NS median: r = 0.73; BS median: r = 0.62). This implied audiovisual STRFs were similarly structured, but with either larger or smaller time-frequency bin values than auditory STRFs for units with positive and negative correlations between auditory STRFs and ΔSTRFs, respectively.

**Figure 11.**
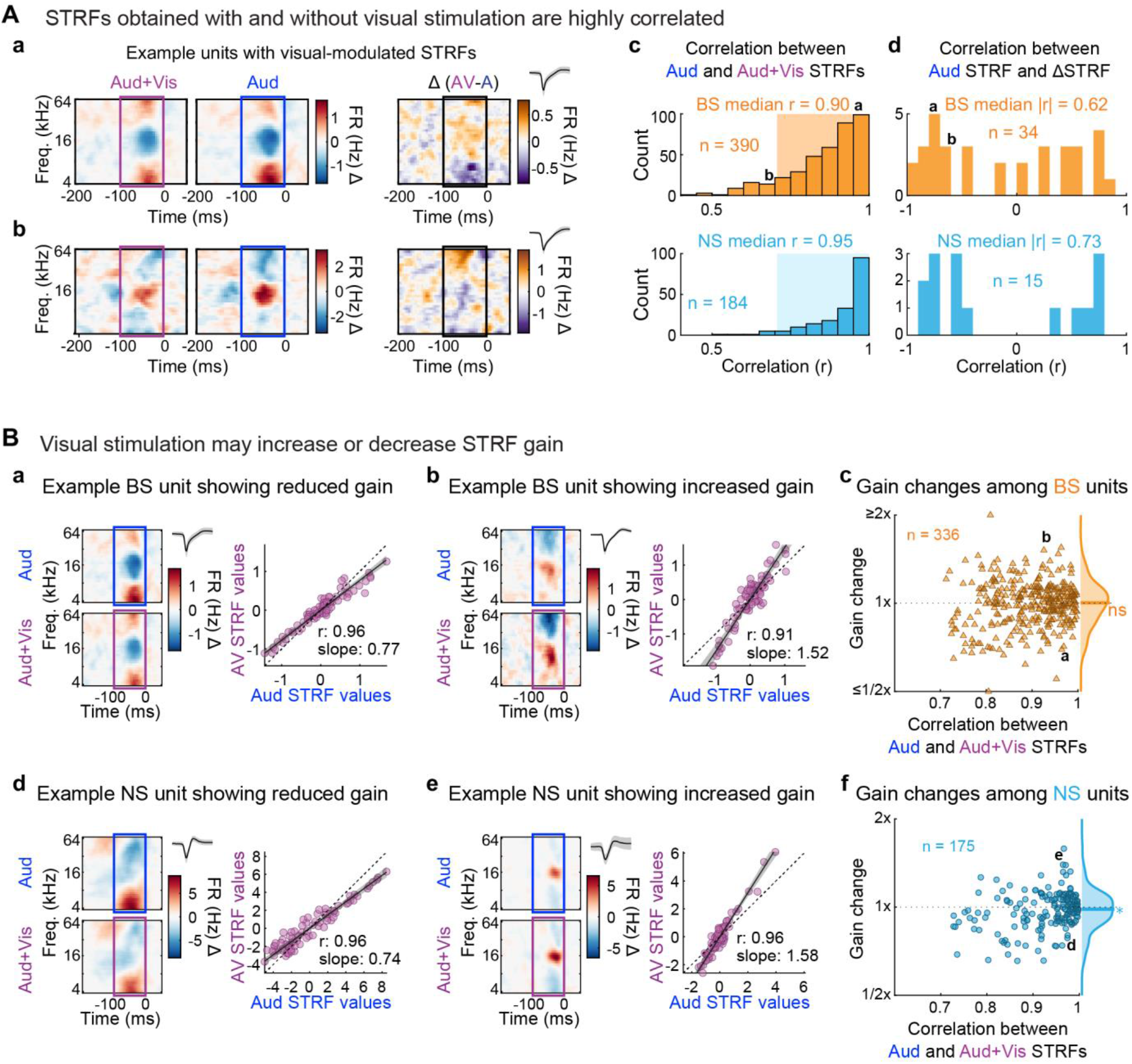
Visual stimulation may change STRF response magnitude but preserves spectrotemporal tuning. (**A**) STRFs obtained with and without visual stimulation are highly correlated. (**a**) Example unit with highly correlated STRFs between conditions. (**b**) Example unit with moderately correlated STRFs between conditions. (**c**) Distributions of STRF correlations between conditions for BS (top) and NS units (bottom), indicating similar spectrotemporal tuning for the majority of units. (**d**) Distributions of correlations between Auditory and ΔSTRFs for each BS (top) and NS units (bottom). ΔSTRFs are typically highly correlated or anti-correlated with Auditory STRFs, suggesting visual stimulation produces global increases or decreases in time-frequency bin values without substantial modifications in STRF structure. (**B**) Visual inputs may alter STRF gain without altering spectrotemporal tuning. (**a**) Example BS unit with gain decrease in the Audiovisual condition. Left: STRFs for each condition with unit waveform (median ± MAD). Right: best fit lines to STRF time-frequency bins from each condition (shading indicates 95% confidence intervals). (**b**) Example BS unit with gain increase in the Audiovisual condition. Subplot organization as in (b). (**c**) Summary of STRF gain changes between conditions as a function of STRF correlations between conditions for BS units. Values above and below 1 (y-axis) indicate higher or lower gain in the Audiovisual condition, respectively. Deviations from 1 were observed even for units with highly correlated STRFs between conditions. Units with increases and decreases in gain were in approximate balance, such that no significant group-level difference from 1 was observed. (**d–e**) Example NS units with decreased and increased gain and in the Audiovisual condition, with subplot organization as in (a–b). (**f**) Summary of STRF gain changes between conditions as a function of STRF correlations between conditions for NS units. Decreases in gain were significantly more common than increases, such that the unit population median was significantly below 1. Wilcoxon signed-rank tests (paired): *p<0.05.

Changes in STRF gain were modeled using standardized major axis regression, which is designed for capturing bivariate relationships with estimation error affecting both variables (Warton et al., 2006). With the auditory and audiovisual conditions plotted on the x and y axes, respectively, positive and negative slopes for the regression model implied that visual stimulation increased and decreased gain, respectively. Slopes were only analyzed for units for which the relationship between conditions could be modeled with high accuracy (r^2^ > 0.5). STRFs were first downsampled by a factor of three in both dimensions to avoid inflating correlations between conditions resulting from the smoothing operation (see Materials and Methods). Example units with gain decreases and increases are shown in **Figure 11B, a–b** for BS units and **Figure 11B, d–e** for NS units. Group data in **Figure 11B, c–f** indicated that gain changes were diverse, showing both increases and decreases. In extreme cases, gain was nearly doubled or halved. However, STRFs were highly correlated between conditions even for units with large gain changes, implying STRF differences were more quantitative than qualitative. As indicated by the y-axis marginal histogram in **Figure 11B, c**, the median gain change was not significantly different from one for BS units (Wilcoxon signed-rank test: p = 0.888), implying units with gain increases and decreases were approximately balanced. A small effect of borderline significance suggesting an overall tendency toward decreased gain (median 0.984) was observed for the NS unit population (Wilcoxon signed-rank test: p = 0.044).

### Visual stimulation may increase or decrease auditory information

The analysis above showed that STRFs for some neurons either increased or decreased in gain on audiovisual trials, i.e., showed greater firing rate deviations from the mean driven rate. This suggested that concurrent visual stimulation may have ‘helped’ or ‘hurt’ encoding of the RDS sound features by individual neurons. However, gain change analysis was restricted to units for which STRFs from each condition were highly correlated (r^2^ > 0.5), which also implied these units had highly STRF reliability (since an unreliable STRF in one condition would not be well correlated with the STRF from the other condition). We further explored the idea that visual stimulation might facilitate or impede encoding of sounds in A1 using an information theoretic approach to quantify visual-mediated changes in information between the auditory stimulus and spike events. This analysis was inclusive of all units with at least 200 driven spikes (to avoid undersampling concerns) regardless of STRF reliability or significance, including potential units with poorly or randomly structured STRFs in one condition or the other. Because previous studies have found multisensory interactions are often dependent upon baseline (unimodal) responses (Allman and Meredith 2007), we further broke down information changes according to baseline information rates in the auditory condition.

As depicted in **Figure 12A, a–c**, similarity between the STRF and stimulus was quantified by convolution, producing a projection value (x) for each time bin used in STRF calculation. The full sample of projection values defined the distribution p(x), which was standardized by its mean and standard deviation. Relatively high and low projections values represent stimulus segments that are similar and dissimilar to the STRF, respectively (examples of each are superimposed over the binned stimulus in **Figure 12A, b**). For auditory responsive units with significant STRFs, spikes are more likely to occur in time bins with high projection values, and least likely to occur in bins with low projection values. This can be demonstrated by identifying the subset of projection values that coincide with spikes, i.e., the distribution p(x|spike). As shown in **Figure 12A, d**, values of p(x|spike) tend to be higher than p(x) for positive standardized projection values, implying increased spike probability for closer matches between the STRF and stimulus. Values of p(x|spike) moreover tend to be lower than p(x) for negative projection values, implying poor matches between the STRF and stimulus result in spiking probability below the mean firing rate. Mutual information is proportional to the ratio of these distributions, as defined by pseudocode in **Figure 12A, d**. Intuitively, p(x) and p(x|spike) become asymptotically equivalent over time for randomly a spiking neuron but diverge to the degree to which a neuron is selectively activated by close stimulus approximations to its receptive field (and suppressed otherwise). Thus, spikes from the example unit in **Figure 12A** are said to be informative about the stimulus because we can infer that the structure of the stimulus is relatively similar to the STRF when the ratio between p(x|spike) and p(x) is high, and relatively dissimilar when the ratio is low.

**Figure 12.**
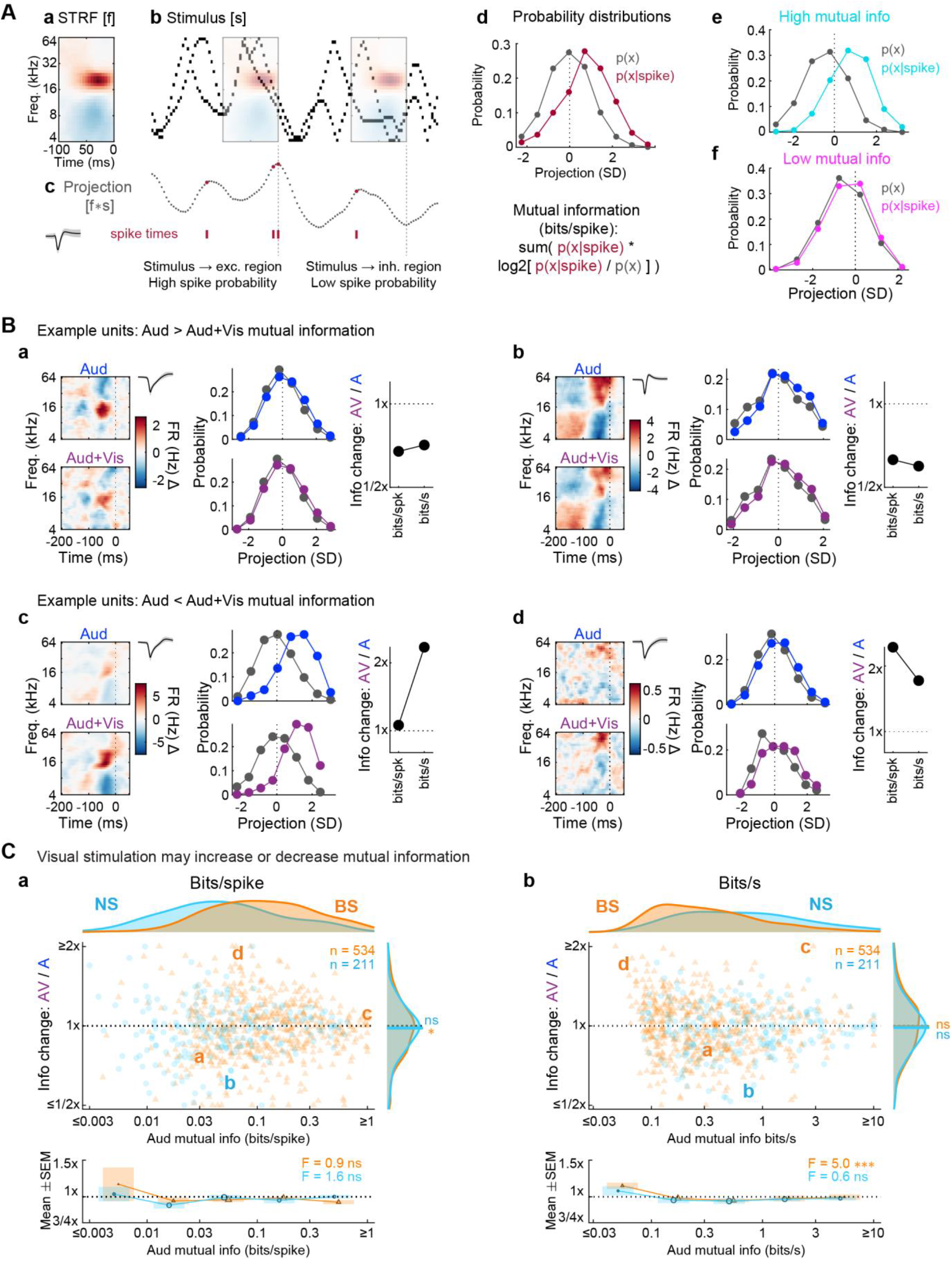
Visual stimulation may increase or decrease information carried about an auditory stimulus. (**A**) Mutual information estimation procedure. (**a**) Example STRF, which serves as filter [f]. (**b**) Example RDS stimulus segment [s]. The overlaid STRFs highlight timepoints at which the RDS frequency vectors intersect excitatory and inhibitory regions of the STRF. (**c**) Projection values (A.U.) for each time point reflecting the convolution of the STRF and stimulus [f*s]. Relatively high and low projection values result from intersections of stimulus energy with excitatory and inhibitory regions of the STRF. Unit waveform shown at left (median ± MAD). (**d**) Distributions of all normalized projection values (gray) and those at time points for which spikes occurred (red). High and low projection values are associated with increases and decreases in spiking probability, respectively. Mutual information between the stimulus (projection value) and response (spike) is estimated from the relationship between these distributions as indicated by the pseudocode below the distributions. (**e–f**) Examples of projection value distributions (hypothetical) which would produce high (**e**) and low (**f**) mutual information values. (**B**) Example units for which visual context decreases and increases mutual information between sound stimuli and spiking responses. (**a–b**) Example units with visual-driven decreases in auditory information. Each example plot includes STRFs (left) and projection value distributions (middle) for each condition, plus an information change summary plot (right) depicting rate changes in both bits/spike and bits/s. Unit waveform shown at to the right of the Auditory STRF (median ± MAD). (**c–d**) Example units with visual-driven increases in information, with subplots organized as in (a–b). (**C**) Summary of changes in mutual information with the addition of visual stimulation, as a function of information in Auditory trials. (**a**) Summary of information changes expressed in bits/spike. Across unit populations, bits/spike tended to be higher for BS units (x-axis marginal). As depicted by the scatter plot (top), information changes were highly heterogeneous across both unit types, with both increases and decreases. Letters correspond to example units in (a–d). As indicated by the marginal histograms for the y-axis, a significant group level bias was observed for BS units only (medians shown by colored lines), with a tendency toward decreased information (Wilcoxon signed-rank tests [paired]: *p<0.05, ns p>0.05). As indicated by binned means below the x-axis of the scatter plot (mean ± SEM), increases and decreases in bits/spike were not significantly dependent upon baseline Auditory information values for either unit type (one-way ANOVA: *p<0.05, **p<0.01, ***p<0.001, ns p>0.05). (**b**) Summary of information changes expressed in bits/s. Across unit populations, bits/s tended to be higher for NS units (x-axis marginal). No significant group-level biases were observed for either unit type (y-axis marginal). As indicated by binned means below the x-axis of the scatter plot (mean ± SEM), increases in bits/s were more likely for BS units with extremely low Auditory information values, with decreases more likely for moderate to high values. No significant relationship was observed for NS units.

A total of 211 NS units and 534 BS units were included in the analysis, reflecting the subset of the full dataset with at least 200 driven spikes and information values greater than zero. Example units with visual-mediated decreases and increases in mutual information are presented in **Figure 12B, a–d**. At the population level, we found that BS units had significantly higher bits per spike than NS units (**Figure 12C, a**; F = 72.17, p < 10^−15^, *η*^2^ = 0.088). However, considering BS units tend to have lower firing rates than NS units, we also analyzed information in terms of bits per second by multiplying the bits/spike value for each unit by its mean firing rate obtained from the sustained response window (spikes/s). The result indicated significantly higher bits/s values for NS units (**Figure 12C, b**; F = 58.33, p < 10^−13^, *η*^2^ = 0.072).

Changes in information rates for individual units were generally consistent with the gain change results reported above. Information rates in the audiovisual condition could be either greater or less than in the auditory condition, for both BS and NS units and for both bits/s and bits/spike. As indicated by y-axis marginals in **Figure 12C, a**, a small but significant decrease in median bits/spike was observed for BS units (Wilcoxon signed-rank test: p = 0.013) but a similar trend for NS units was not significant (Wilcoxon signed-rank test: p = 0.072). Changes in bits/spike did not significantly depend upon baseline auditory information rates for either unit type (BS units: F = 0.874, p = 0.479, *η*^2^ = 0.007; NS units: F = 1.61, p = 0.174, *η*^2^ = 0.030). Population level decreases in bits/s were non-significant for both unit types (Wilcoxon signed-rank tests: BS units: p = 0.385; NS units: p = 0.067). For BS units, changes in bits/spike depended significantly upon baseline auditory information rates (F = 5.01, p < 10^−3^, *η*^2^ = 0.037), such that units with very low auditory information (<0.1 bits/s) on average showed increases in information transfer with visual stimulation, whereas information for most other units decreased. A similar relationship was not observed for NS units was not significant (F = 0.613, p = 0.654, *η*^2^ = 0.012).

Considered together, gain and information change analyses produced three insights about the influence of visual context on sound encoding in A1. First, effects are highly heterogeneous at the individual unit level, including units with both increases and decreases in gain and information. Second, population level changes in gain and information in the audiovisual condition are either subtle or non-significant, and generally reflect decreases relative to the auditory condition. Third, whether visual context facilitates or interferes with sound encoding may partially depend on baseline auditory responsiveness.

## Discussion

Perception is inherently multisensory under natural conditions. Environmental events are often encoded by more than one sensory modality, such as correlated visual and auditory cues supporting spatial perception and conspecific communication (Sugihara et al., 2006; Allman and Meredith 2007; Bigelow and Poremba, 2016). Even unisensory events must be encoded within the context of uncorrelated sensory processing by other modalities. Growing recognition of the ubiquity of such phenomena has resulted in increased attention to multisensory context within the sensory physiology community, which has traditionally been dominated by unimodal paradigms. One of the most important insights from these efforts is that multisensory interaction is pervasive throughout cortex, including direct anatomical connections and functional interaction between primary sensory cortices (Ghazanfar and Schroeder, 2006; Bizley et al., 2007; Banks et al., 2011; Iurilli et al., 2012; Bizley et al., 2016; McClure and Polack 2019). For instance, a recent study from our lab found that a subset of neurons in awake mouse A1 were responsive to visual flash stimuli alone, and that these neurons were concentrated in the infragranular layers (Morrill and Hasenstaub, 2018). These outcomes were replicated and extended in the present study using the CMN visual stimulus (**Figure 2**), which also confirmed that both putative inhibitory (NS) and excitatory neurons (BS) can be visually responsive. Of these, all but a few were also significantly responsive to the RDS sound stimuli (**Figures 3, 5– 6**), suggesting purely visual neurons in A1 are very rare.

The current study further identified neurons in A1 for which responses to sound were modulated by visual stimulation (**Figures 7–12**). A major question motivating the present study was whether there was close overlap between units with such visual-modulated responses and units responsive to visual stimulation alone. If so, the expected depth distribution of visual-modulated responses to sound would be similarly concentrated in the deep cortical layers, as was recently observed in a human imaging study (Gau et al., 2020). We obtained only mixed support for this hypothesis. Fewer than half of neurons with visual-modulated sound responses showed significant responses to visual stimulation alone. These outcomes replicate the findings of previous studies that unimodal visual responses are neither necessary nor sufficient for visual-modulated responses to sound. Consistent with these observations, the depth distributions of units with visual-modulated sound responses and units with unimodal visual responses were, at best, only weakly similar. For instance, in parallel with unimodal visual responses, we found that visual influences on sound sustained firing responses depended on cortical depth, with the majority of significant effects in the deepest cortical bin (**Figure 7**). However, similar depth dependencies were not observed for modulated onset responses or STRFs. Together, these outcomes support a model of audiovisual integration wherein visual projections to infragranular A1 drive responses to visual stimulation alone, after which such activity may modulate sound-evoked responses by propagating throughout cortical layers.

By delivering segmented RDS stimuli separated by intertrial intervals, we were able to estimate time-averaged firing rate changes from baseline reflecting both transient onset responses and sustained responses throughout the sound period. Moreover, averaging the binned stimulus values within a window preceding each spike event time enabled estimation of STRFs, which are sensitive to both spike count and timing with respect to the stimulus. Capturing each response type turned out to be important both for characterizing responses to the sounds themselves and for the dependence of these responses on visual context. Notably, a double dissociation was observed for onset and sustained firing responses, wherein much stronger baseline deviation effects were observed for onset responses on auditory trials (**Figure 3**), but much stronger visual modulation effects were observed for sustained responses on audiovisual trials (**Figure 7**). We further found that neither firing rate response was necessarily present in units with significantly reliable STRFs (**Figure 4**). Similarly, only partial overlap was observed among units with visual-modulated sustained firing rate responses and units with visually modulated STRFs, even though the spikes entering each analysis were identical (**Figure 10**). Collectively, these outcomes reinforce previous studies concluding that spike rate and timing changes in A1 carry non-redundant information about the stimulus (Brugge and Merzenich, 1973; deCharms and Merzenich 1996; Recanzone, 2000; Lu et al., 2001; Wang et al., 2005; Malone et al., 2010; Malone et al., 2015; Insanally et al., 2019; Liu et al., 2019).

An important caveat regarding onset and sustained firing rate responses in the current study is that they were elicited by complex, non-repeating stimuli. This aspect of our study design has at least two implications for the results. First, averaged spike rates within any given time bin, and across full analysis windows, reflect mixed responses to diverse spectrotemporal features produced by random draws from the RDS stimulus distribution. Thus, they likely reflect a combination of weak and strong excitatory responses, as well as suppressed responses resulting from intersections of spectral energy with receptive field inhibitory sidebands, refractory periods, and from simultaneous and forward masking effects the stimulus was designed to produce (Gourévitch et al., 2015). By contrast, binned spike averages elicited by a repeating RDS stimulus segment would reflect specific spectrotemporal features that drive or inhibit spiking activity. A neuron without a significant change in averaged sustained firing in a non-repeating stimulus paradigm may thus exhibit regular temporal patterns of excitation and suppression in a repeating stimulus paradigm. Second, onset and sustained responses observed in the current study are not directly comparable to many previous studies examining diverse firing rate responses in A1 (onset, sustained, offset), the majority of which presented repeated tone pips, often at the best frequency of the neuron (Brugge and Merzenich, 1973; Recanzone, 2000; Wang et al., 2005; Malone et al., 2015; Liu et al., 2019). Thus, it cannot be assumed that onset and sustained responses observed in previous studies would necessarily be subject to similar asymmetries in terms of visual context modulation. The diverse averaged firing rate response types observed in the present study are thus qualitatively different from previous results obtained with repeated tone pips, but nonetheless underscore the same conclusion that different firing rate responses contain unique information about the stimulus and are not equivalently sensitive to sensory and other contextual variables.

Previous studies of multisensory integration across multiple modalities in a wide range of cortical and subcortical stations have reported that stimulus encoding may be either facilitated or impeded by the addition of a second modality (Perrault et al., 2003; Romanski, 2007; Stein and Stanford 2008; Sugihara et al., 2010; Kobayasi and Riquimaroux, 2013). Similar outcomes have been obtained with both correlated and uncorrelated bimodal stimulation (Dahl et al., 2010), with several studies suggesting benefits are more common with correlated stimulation (Diehl and Romanski, 2014; Meijer et al., 2017). Costs are thought to possibly reflect competition by each modality over limited spike resources within a given neuron. For example, in a neuron responsive to two modalities, a subset of the spikes otherwise under the control of a single modality may instead be driven by a second modality during bimodal stimulation, thus potentially decreasing the mutual information between spiking and the stimulus of the first modality. In correlated bimodal stimulus paradigms, such as natural audiovisual speech, encoding benefits of bimodal stimulation likely reflect synergistic interaction of temporally and/or spatially coincident feature selectivity within each respective modality. In uncorrelated bimodal stimulus paradigms, including the present study, improved encoding with the addition of an uncorrelated signal from another modality may reflect stochastic resonance, in which sensitivity to a weakly detectable stimulus may be increased even by random noise within or between modalities (Wiesenfeld and Moss, 1995; Shu et al. 2003; Hasenstaub et al. 2005; Crosse et al., 2016; Malone et al., 2017). Consistent with this possibility, some evidence obtained in the present study suggested that auditory information ‘costs’ and ‘benefits’ produced by visual stimulation were non-randomly distributed with respect to baseline information rates obtained with auditory stimulation alone. Specifically, information increases for BS units (bits/s) were most likely for units with the lowest auditory information rates, suggesting weak and inconsistent responses to sound features may be strengthened and regularized by the additional visual stimulus. These and similar findings by previous studies establish neural phenomena parallel to psychophysical experiments reporting significantly improved detection of weakly perceptible events following the addition of both correlated and uncorrelated stimulation within a second modality (Ward et al., 2010; Huang et al., 2017; Gleiss and Kayser, 2014; Krauss et al., 2018).

## Acknowledgements

This work was supported by National Institutes of Health grants F32DC016846 to J.B and R01DC014101, R01NS116598, R01MH122478, and R01EY025174 to A.R.H., the National Science Foundation GRFP to R.J.M., the Klingenstein Foundation to A.R.H., Hearing Research Inc. to A.R.H., PBBR Breakthrough Fund to A.R.H., and the Coleman Memorial Fund to A.R.H.

